# Genetic determinants of distinct CD8^+^ α/β-TCR repertoires in the genus *Mus*

**DOI:** 10.1101/2024.09.05.611437

**Authors:** Moritz Peters, Volker Soltys, Dingwen Su, Marek Kučka, Yingguang Frank Chan

## Abstract

The adaptive immune system’s efficacy relies on the diversity of T cell receptors and the ability to distinguish between self and foreign antigens. Analysis of the paired heterodimeric αβ-TCR chains of individual T cells requires single-cell resolution, but existing single-cell approaches offer limited coverage of the vast TCR repertoire diversity. Here we introduce CITR-seq, a novel, instrument-free, high-throughput method for single-cell TCR sequencing with >88% αβ-TCR pairing precision. We analyzed the TCR repertoires of CD8^+^ T cells originated from 32 inbred mice using CITR-seq, comprising four evolutionary divergent sister species and their F1 hybrids. Overall, we identified more than 5 million confidently paired TCRs. We found that V(D)J gene usage patterns are highly specific to the genotype and that Vβ-gene usage is strongly impacted by thymic selection. Using F1 hybrids, we show that differences in gene segment usage across species are likely caused by *cis*-acting factors prior to thymic selection, which imposed strong allelic biases. At the greatest divergence, this led to increased rates of TCR depletion through rejection of particular Vβ-genes. TCR repertoire overlap analysis across all mice revealed that sharing of identical paired CDR3 amino acid motifs is four times more frequent than predicted by random pairing of TCRα and TCRβ chains, with significantly increased sharing rates among related individuals. Collectively, we show that beyond the stochastic nature of TCR repertoire generation, genetic factors contribute significantly to the shape of an individual’s repertoire.

## Introduction

Adaptive immunity relies on the recognition of antigens presented on the surface of virtually all nucleated cells through class I and II major histocompatibility complexes (MHC). These MHC-bound antigens are recognized by T cell receptors (TCRs) expressed on the surface of T cells, which collectively possess the remarkable ability to discriminate between antigens of “self” and “foreign” origin. This is critical for preserving self-tolerance, thereby preventing autoimmunity, while also enabling the identification of pathogen-infected or malignant cells to initiate an immune response [1]. The nature of an immune response depends largely on whether an antigen is presented via class I or class II MHC complexes, which are targeted by CD8^+^ cytotoxic T cells [2] or CD4^+^ helper T cells [3], respectively. The former can induce apoptosis in targeted cells while the latter can trigger secondary immune cascades involving B-lymphocytes and cells of the innate immune system. In both cases, TCRs are the key molecules that mediate signaling and enable a broad spectrum of immune responses.

TCRs are primarily composed of two heterodimeric chains, TCRα and TCRβ, both of which arise from somatic rearrangements of gene segments during T cell development. This rearrangement process, known as V(D)J-recombination, generates diversity through joining variable (V), diversity (D, exclusive to TCRβ) and joining (J) gene segments to a constant region, thereby generating a unique TCR receptor in each individual T cell [4]. Additional diversity is introduced through nucleotide insertions and deletions at each junction during V(D)J-recombination [5]. In the expressed TCRα and TCRβ chains, the resulting highly polymorphic junctional region is situated in closest proximity to antigens presented by MHCs to serve as a binding pocket [6] and is termed complementarity-determining region 3 (CDR3). The other CDRs, 1 and 2, constitute germline encoded regions within the V-segments of TCR chains and are believed to primarily facilitate TCR-MHC binding and are less relevant for antigen recognition [7, 8] (**Fig. 1A**).

**Figure 1:**
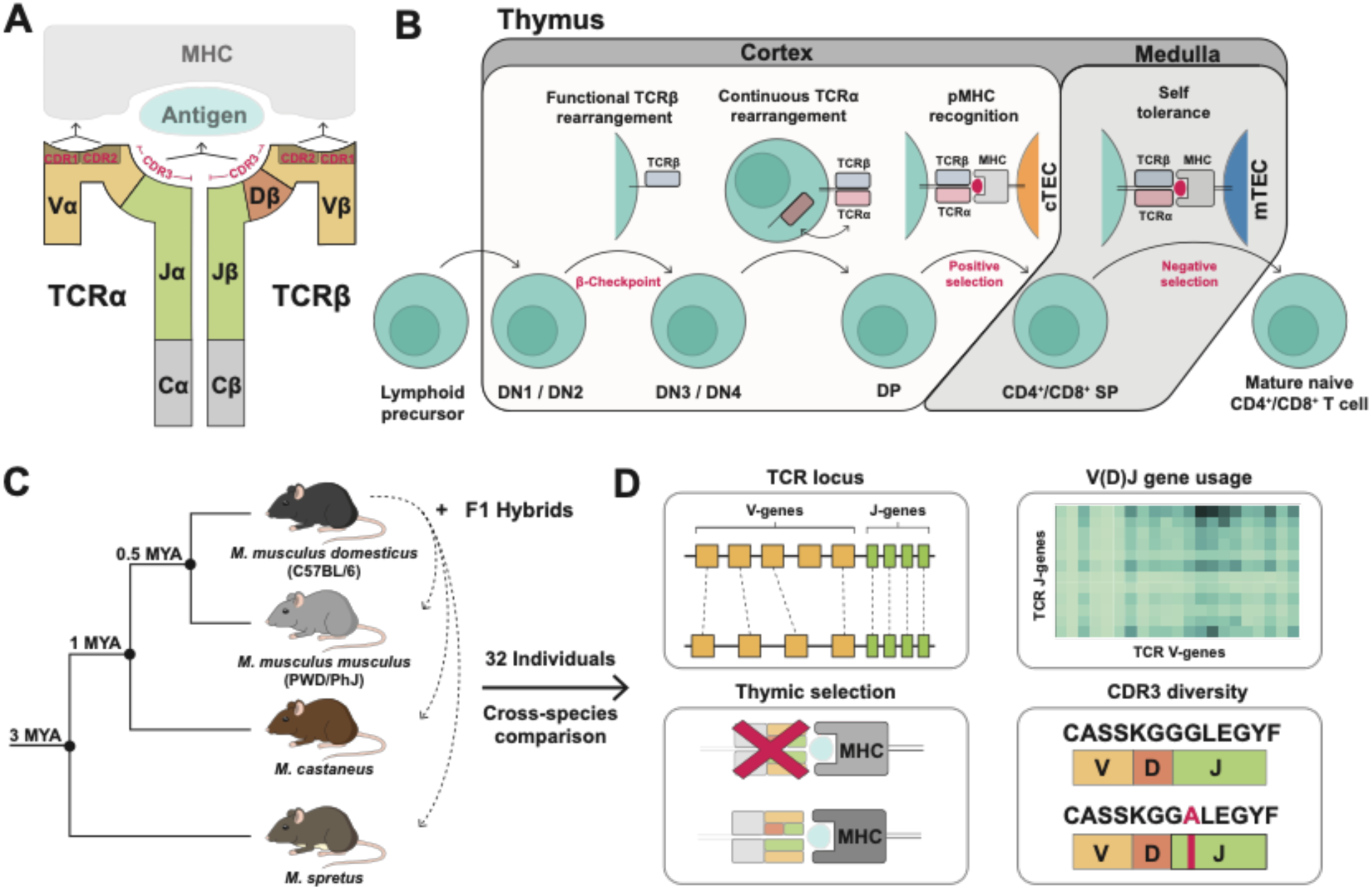
Introduction to T cell receptors and overview of the study design. **(A)** Heterodimeric αβ-TCR consisting of a V-, D- (exclusive to TCRβ) and J-gene alongside the constant (C) region. The junctional region of V(D)J genes marks the CDR3 sequence that is in the closest proximity to the antigen in the TCR-MHC complex. CDR1 and CDR2 are germline-encoded sequences of V-genes that contribute to TCR-MHC binding. **(B)** T cell maturation in the thymus. Intra-thymic T cells are classified by the expression of the lineage-markers CD4 and CD8 (double negative: DN, double positive: DP and single positive: SP). T cells that successfully rearranged a functional TCRβ chain can pass the β-checkpoint. Subsequent continuous rearrangements of the TCRα chain leads to transition to the DP stage. T cells with a fully assembled αβ-TCR that is capable of binding self-MHCs on the surface of cTECs survive positive selection. Selected T cells migrate to the medulla and undergo negative selection, during which T cells that strongly bind self-MHCs on the surface of mTECs are rejected. T cells that survive both selection steps are released from the thymus. **(C)** Phylogenetic tree showing the evolutionary divergence of inbred mouse species used in this study. **(D)** The different aspects of TCR generation and selection analyzed in the course of this study.

The antigen specificity of each rearranged TCR as well as its affinity to MHCs is evaluated in a key multi-step process called thymic selection. It takes place during T cell maturation which is generally classified by the intra-thymic progression from the CD8/CD4 double negative (DN) stages of lymphoid precursors to the single positive stage (SP) of mature T cells. The selection process is initiated at the so called β-checkpoint [9], at which the successful rearrangement of an in-frame TCRβ chain is controlled. Afterwards, the fully assembled αβ-TCR is tested for its affinity to MHCs during positive selection and its specificity to presented self-antigens during negative selection. Both processes are chronologically and spatially separated: Positive selection occurs in the cortex through interactions with cortical thymic epithelial cells (cTECs), whereas the subsequent negative selection occurs in the medulla through interactions with medullary thymic epithelial cells (mTECs) (reviewed here [10]). Overall, only about 5% of T cell precursors survive thymic selection of their TCRs by demonstrating adequate affinity to MHC complexes while simultaneously exhibiting tolerance towards the broad spectrum of presented self-antigens and thus thymic selection significantly decreases the diversity in the TCR repertoire (**Fig. 1B**).

In our current understanding, nucleotide insertions and deletions, V(D)J-segment usage and αβ-TCR pairing are mostly seen as stochastic events that give rise to highly unique and dynamic TCR repertoires within and across individuals [11]. However, recent work increasingly suggests that the diversity of TCR repertoires also relies on genetically encoded differences across individuals [12-15]. For example, V-segment usage in identical twins exhibits much greater similarity compared to unrelated individuals [16]. Notably, this provides evidence that genetics may operate at two different levels: V(D)J-recombination as well as thymic selection, as indicated by the fact that identical twins also share the same set of MHC class I and II (also known as human leukocyte antigen, or HLA) alleles. This observation, and the multi-level genetic determinants that collectively shape TCR diversity, are the focus of at least two debates: one concerning the existence of a co-evolutionary feedback process between TCR and MHC binding [17-21], and the other on whether HLA heterozygosity is evolutionarily optimal due to the presentation of a broader immunopeptidome [22, 23], or alternatively, deleterious due to a high frequency of presented self-peptides, leading to increased depletion of autoreactive TCRs [24, 25]. The extreme diversity of both binding partners, TCR and MHC, makes answering these questions extremely challenging. By contrast, panels of inbred mouse lines spanning within- and across-species diversity, along with their F1 hybrids, provide a tractable setup to address the question regarding the role of MHC/HLA alleles in shaping repertoire diversity during thymic selection.

Estimates on the theoretical αβ-TCR diversity vary greatly by species as well as methodology. Initial theoretical estimates on αβ-TCR diversity were approximately 10^15^ in mice [4] and 10^18^ [26] – 10^20^ [27] in humans. More recent calculations now greatly exceed those estimates and range up to 10^61^ [28]. However this needs context: the number of theoretical αβ-TCRs vastly outnumbers the actually realized TCRs in the repertoire of an individual, primarily because the number of present T cells of an individual (10^12^ [29] in humans and 10^8^ [30] in mice) is several orders of magnitude smaller at any given time. Interestingly, despite the great difference in number of total T cells across different species (e.g. more than 1000x more T cells in humans than in mice), the diversity within the realized TCR repertoire has shown to be much more similar across species [30, 31]. This observation gave rise to the idea of a minimally required repertoire size defined as a functional unit of the “protecton” which is simply multiplied in species with larger numbers of total T cells [32]. Experimental validation of TCR repertoire diversity estimates still suffers from the limitations of current methodologies. While bulk assays can now feasibly analyze entire repertoires across many individuals [33-35], they leave out the critical pairing between TCRα and TCRβ chains within individual cells. Paired TCR analysis requires molecular barcoding of single cells, but existing methods often rely on pre-existing single-cell workflows, restricting analysis to thousands of T cells rather than entire repertoires [36-38]. These protocols typically utilize fluorescence-activated cell sorting (FACS), or microfluidic platforms to isolate individual cells and therefore require specialized equipment. The recent development of SPLiT-seq [39] has expanded the scope of single-cell whole transcriptome experiments to up to 10^6^ cells per experiment by using combinatorial indexing to molecularly barcode each individual cell. Despite this increase in throughput, the associated sequencing cost still substantially limits the feasibility of these methods for assessing large TCR repertoires, especially across multiple individuals.

Here, we investigate repertoires of paired αβ-TCRs from cytotoxic CD8^+^ T cells by developing a targeted TCR sequencing protocol called CITR-seq (**C**ombinatorial **I**ndexing **T** cell **R**eceptor sequencing) to analyze TCR repertoires at low cost and large scale. We apply CITR-seq to 32 individual mice from 4 distinct inbred sister species (C57BL/6J, CAST/EiJ, PWD/PhJ, SPRET/EiJ, abbreviated as BL6, CAST, PWD and SPRET, respectively) and their F1 hybrids with BL6, spanning an evolutionary divergence of approximately 3 million years [40-42] (**Fig. 1C**). The diverged but controlled genetic backgrounds provide a unique opportunity to determine the respective impact of TCR locus structure, the V, D and J gene segment usage frequency, TCR/MHC allele co-evolution via thymic selection and ultimately the joint effects on CDR3 diversity (**Fig. 1D**).

## Results

### CITR-seq design and validation

To generate TCR repertoires we built on SPLiT-seq [39] to develop CITR-seq and modify the approach to generate RNA-based targeted paired αβ-TCR libraries (**Fig. 2A**). We first isolated CD8^+^ T cells from spleens of 10-week-old mice by using anti-CD8 magnetic beads and subsequent purification by FACS (**Fig. S1A**). Purified CD8^+^ T cells were either used directly or were transferred to anti-CD3 and anti-CD28 coated tissue culture plates for a 20-hour activation period before the library preparation (see **Suppl. Table 1** for detailed sample list; note that we limited the activation to 20 hours to avoid cell doubling). Primary or activated T cells were fixed and permeabilized and a set of barcoded TCRα and TCRβ constant-region primers was used to perform *in situ* reverse transcription (RT) inside individual cells (see **Suppl. Table 2** for a list of all primers and barcoding DNA-oligos). All RT-primers contain a unique molecular identifier (UMI) and a ligation overhang for single-cell barcoding. Here, cells are distributed in two split-and-pool cycles across two 96-well plates, such that the RT-primer overhangs are ligated to oligos carrying barcode segments. The split-and-pool approach allow all reactions to be performed in bulk, while giving each cell an effectively unique barcode (calculated barcode collision rate: 1.67%; **see methods**). Afterwards up to 10,000 cells are merged into sub-samples and reverse crosslinking is done to make the barcoded cDNA accessible for amplification. Second strand synthesis is done in a multiplex-PCR setup using 54 primers targeting the 5’ ends of TCR V-segments. In a final index-PCR another DNA barcode is added to each sub-sample, which expands the total barcoding space to up to 28 million unique cellular barcodes. Subsamples can then be pooled to generate the final sequencing-ready library which is compatible with standard Illumina workflows and can be further multiplexed with other sequencing libraries (**Fig. 2B**). This process is cost efficient (**see methods**) and does not require specialized instrumentation.

**Figure 2:**
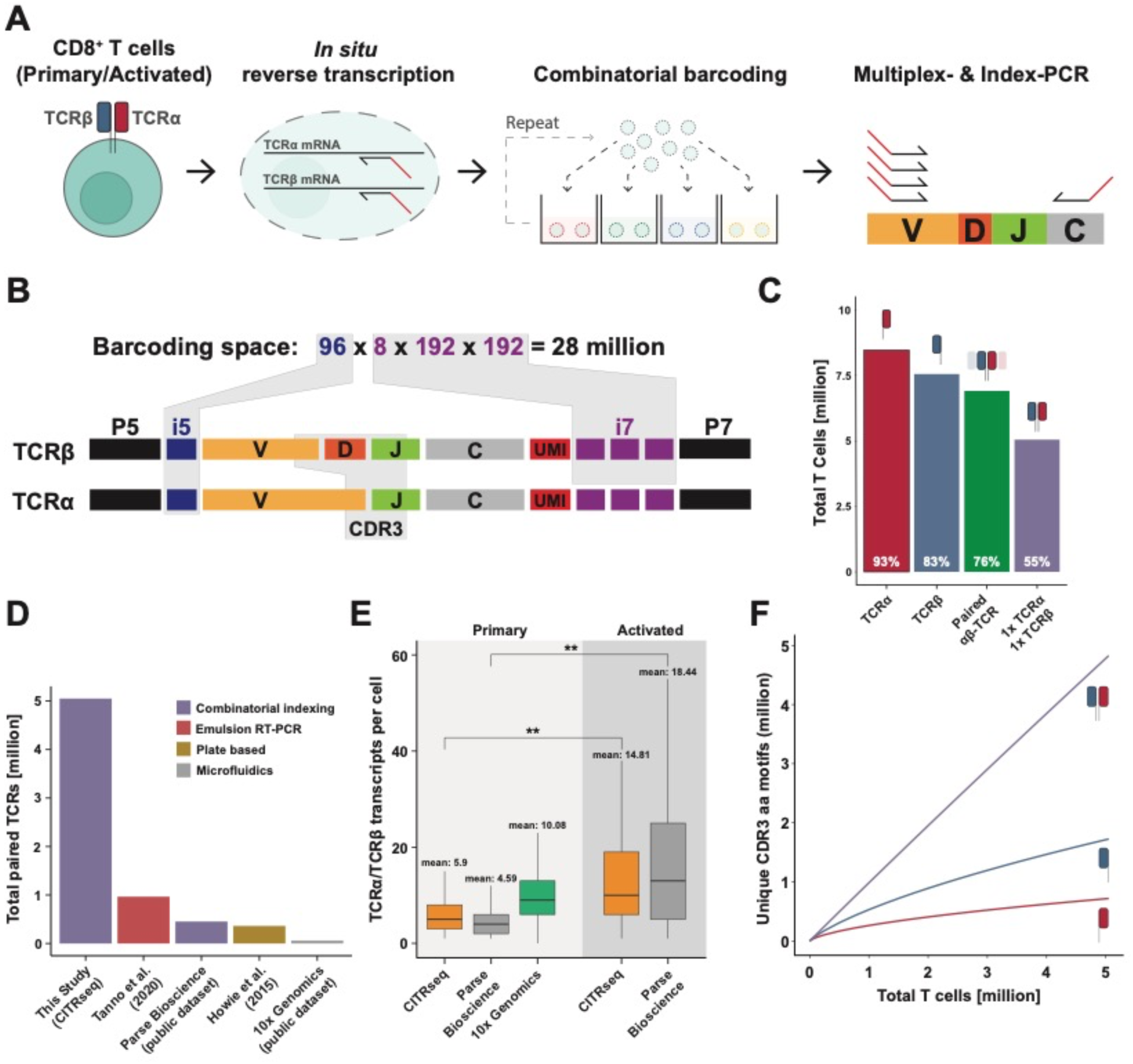
CITR-seq allows for the analysis of millions of confidently paired αβ-TCRs. **(A)** Workflow for generating paired αβ-TCR sequencing libraries using CITR-seq. Isolated T cells are fixed and permeabilized. TCRα and TCRβ are *in-situ* reverse transcribed using barcoded primers targeting both TCR constant regions. Afterwards, T cells are distributed across 96-well plates and well-specific barcodes are ligated to the cDNA. This process is repeated once by pooling all T cells and redistributing them to a second set of barcoding plates. T cells are then pooled again and split into sub-samples before reverse crosslinking. Second strand cDNA is generated in a multiplex-PCR with primers targeting the 5’ region of Vα- and Vβ-genes. In a final Index-PCR a fourth barcode is added. **(B)** Barcode and sequencing adapter structure in CITR-seq libraries. The different combinations of all four barcodes provide a barcoding space of more than 28 million possible barcodes. Sequencing reads fully cover CDR3α and CDR3β sequences. **(C)** Pairing rate across all CITR-seq samples in this study. Fraction of the 9.113.392 total T cells that were assigned to a TCRα (93%, red), a TCRβ (83%, blue), at least one TCRα and TCRβ (76%, green) or exactly one TCRα and TCRβ chain (55%, violet). **(D)** Total number of paired αβ-TCRs analyzed in different studies and publicly available datasets generated with different methods. Emulsion RT-PCT [14], PairSeq (plate-based) [43], two publicly available datasets generated with combinatorial indexing (Parse Bioscience) [44] and microfluidics (10x Genomics) [45]. **(E)** Mean number of TCRα and TCRβ transcripts (UMIs) per cell-barcode in primary and activated T cells in CITR-seq samples (primary and 20h activated mouse T cells), across all 10x Genomics Single-Cell Immune Profiling libraries (primary mouse T cells) and in two publicly available datasets from Parse Bioscience (primary and 72h activated human T cells) [44, 46]. **(F)** Total number of unique CDR3α, CDR3β or paired CDR3αβ amino acid motifs relative to the number of T cells across all 32 CITR-seq samples.

Using CITR-seq, we profiled TCR repertoires from a total of 9,113,392 CD8^+^ T cells (hereafter referred to as “T cells”) across all 32 individuals. Paired TCRα and TCRβ chains were successfully recovered in 75.8% of T cells, 55.4% of which carried exactly one α- and β-chain (**Fig. 2C**). To the best of our knowledge, this dataset of 5,049,334 singly α/β- paired T cells represents the largest set of paired TCRs analyzed in a single study to date (**Fig. 2D**). To assess pairing precision, we determined the rate of repeated observations of identical Vβ-Jβ-CDR3β and Vα-Jα-CDR3α mates (or “clonotypes”) in a sample of 150,000 T cells that underwent clonal expansion for 72h in tissue culture. Clonal expansion through prolonged tissue culture allowed us to enrich for cells carrying the same α/β-chain pairing, the recovery of which would have been unlikely under our standard protocol. TCRβ chains that were observed at least twice in this repertoire were seen with identical TCRα chains in 88% of cells, thus representing T cells with identical clonotypes. This rate marks the lower-bound pairing precision, since with 150,000 T cells, we expect a low, but non-negligible chance of recovering the same Vβ-Jβ-CDR3β chain from two non-clonal T cells which should therefore pair with a different TCRα chain. In agreement with this high pairing precision, we observed few cells with more than two TCRα (1.4%) or TCRβ (0.5%) chains across all 32 CITR-seq samples, which is biologically implausible because of the presence of just two alleles for each chain in each T cell (**Fig. S2A**).

We then compared transcripts (UMIs) per cell counts at saturating read coverage (mean reads/cell: 184.57; **Fig. S2B**) in activated and primary T cells in CITR-seq. Activated T cells had a significantly higher UMI/cell count (14.81) compared to primary T cells (5.9, pairwise t-test; *P* < 0.01; **Fig. 2E**). We compared these values to two publicly available human TCR sequencing datasets (Parse Bioscience; **see methods**), in which 72h activated T cells also showed significantly higher average UMI per cell count (18.44 UMIs/cell) than primary T cells (4.69 UMIs/cell). As a further benchmarking effort, we generated complementary datasets for each of the four inbred mouse species from primary T cells using the Chromium Next GEM Single Cell 5’ platform by 10x Genomics (**see methods**). For these, we recovered 10.1 UMIs/cell on average across samples (**Fig. 2E**).

We evaluated whether activation of T cells biases the recovered TCR repertoire by comparing V-J usage (discussed below) and clonal abundance in samples of primary and activated T cells, each down-sampled to 150,000 cells. To do so, we first compared the frequency of multiple observations of identical TCRα and TCRβ as well as full clonotypes (defined by identical V+J+CDR3 amino acid motif) across cells (**Fig. S2C**). In TCR repertoires of primary T cells and T cells that were activated for 20h, most αβ-TCR pairs were exclusive to a single cell (93% and 96.7% respectively). This is in contrast to αβ-TCR pairs in TCR repertoires of cells that were activated for 72 hours, in which less than half (48%) of pairs are exclusive to a single cell with all other pairs being observed multiple times. We therefore conclude, that in agreement with previous studies [79] the 20h activation protocol did exclude clonal expansion of T cells.

### Validation of CITR-seq against 10x Genomics commercial platform

To validate complete coverage of all the functional V/J genes, we compared V-J gene usage frequencies from data generated using CITR-seq and 10x Genomics Immune Profiling. We find high correlation of V-J usage frequencies (Pearson: BL6 r = 0.91) across both methods (**Fig. S3C**). Additionally, the highly correlated V-J usage frequencies provide further evidence for the unbiased repertoire representation of 20h activated (CITR-seq) compared to primary (10x Genomics Immune Profiling) T cells.

To evaluate the coverage of cross-species repertoires, we analyzed CDR3 amino acid motif diversity in α- and β-chains both individually and jointly (**Fig. 2F**). In the 5,049,334 T cells from across all 32 samples, we detect 719,976 (14.26%) unique TCRα and 1,725,631 (34.18%) unique TCRβ chains. If analyzed jointly, 95.6% of these (4,826,991) represent unique αβ-TCR pairs. In contrast, in 9,445 paired T cells in our Chromium Next GEM Single Cell 5’ datasets, we found 85% and 94.9% unique TCRα and TCRβ chains, respectively (n = 8,021 and 8,963), and nearly all (98.6%, or 9,313 T cells) represent unique αβ-TCR pairs (**Fig. S2D**). Taken together, we interpret this data to show the remarkable diversity, especially across paired TCRs: even with the throughput of CITR-seq at 5 million cells, we were not close to sampling T cell clonotypes to saturation, let alone using much more limited platforms. This further emphasizes the need for high-throughput methods to gain reasonable insight into the diversity of TCR repertoires.

### Distinct V-J usage patterns across mouse species

To compare V-J segment usage across the different mouse species, we first constructed species-specific V(D)J-segment references (**see methods**). Across all samples, mapping against the corresponding species-specific reference showed a slight increase in the total number of successfully aligned sequencing reads (PWD: +0.07%, CAST: +0.07% and SPRET: + 0.1% total reads; **Fig. S3A**) and per segment alignment scores (data not shown), relative to mapping against an mm10-based V(D)J reference provided in the MiXCR software [47]. Local alignment of TCR loci to the mm10 reference genome (GRCm38/mm10) revealed one-to-one orthology in Vβ-, Jβ- and Jα-segments. In contrast, we found extensive rearrangements, including inversions and gene-cluster triplications in the Vα cluster between the four mouse species (**Fig S3B**). For instance, the central region of the Vα cluster (Vα gene families 3-15) is expanded in BL6, PWD and SPRET relative to the CAST Vα cluster. This results in a Vα locus size reduction of ∼0.6 Mb and approximately 70 fewer Vα genes in CAST. In a given species, sequence identity across Vα paralogues is extremely high (e.g., in BL6 *Trav11* and *Trav11D* are 100% identical on the nucleotide level; for details see [48]). For this reason, and to properly handle multiple Vα read mapping, we grouped Vα genes into their respective gene families for cross-species comparison.

We compared the mean of intra-species V/J gene segment usage across BL6, PWD, CAST and SPRET mice (**Fig. 3A**). Consistent with previous studies [33, 49], V/J segment usage within an individual typically spanned several orders of magnitude in both TCRα and TCRβ chains. For example, in BL6 TCR repertoires *Trbv12-1* and *Trbj2-7* were found in 2.37% of likely productive, in-frame, TCRβ chains, while the combination of *Trbv21* and *Trbj1-5* was only present in 0.0003% of in-frame TCRβ chains. Across species, segment usage frequencies are broadly similar, with some notable and extreme exceptions, mostly in Vβ-Segments (e.g. *Trbv13-2* is used in 7.75% of BL6 TCRβ chains compared to only 0.14% in SPRET). Consistent with previous studies [50] we observe decreased pairing of proximal Vα and distal Jα as well as distal Vα and proximal Jα segments. The only exception to this rule is CAST, where we observed significantly higher usage frequencies of the most distal Vα gene (*Trav1* and T*rav2*; chi-squared test; ** *P* < 0.01; **Fig S3D**). In laboratory mice, it has been shown that TCRα V-J recombination proceeds progressively from 3’ (proximal) Vα genes towards more 5’ (distal) Vα genes [51]. We therefore hypothesize, that the significantly higher usage of distal Vα genes in CAST is linked to its contracted Vα locus. As a general trend, we found that the average variance in V-segments usage frequency (var(Vβ) = 12.44, var(Vα) = 5.95) was higher compared to the average variance in J-segment usage frequency (var(Jβ) = 4.91, var(Jα) = 0.48) when compared across all species. This indicates that species-specific differences in V(D)J usage mostly arise from biases in V-segment usage rather than J-segment usage.

**Figure 3:**
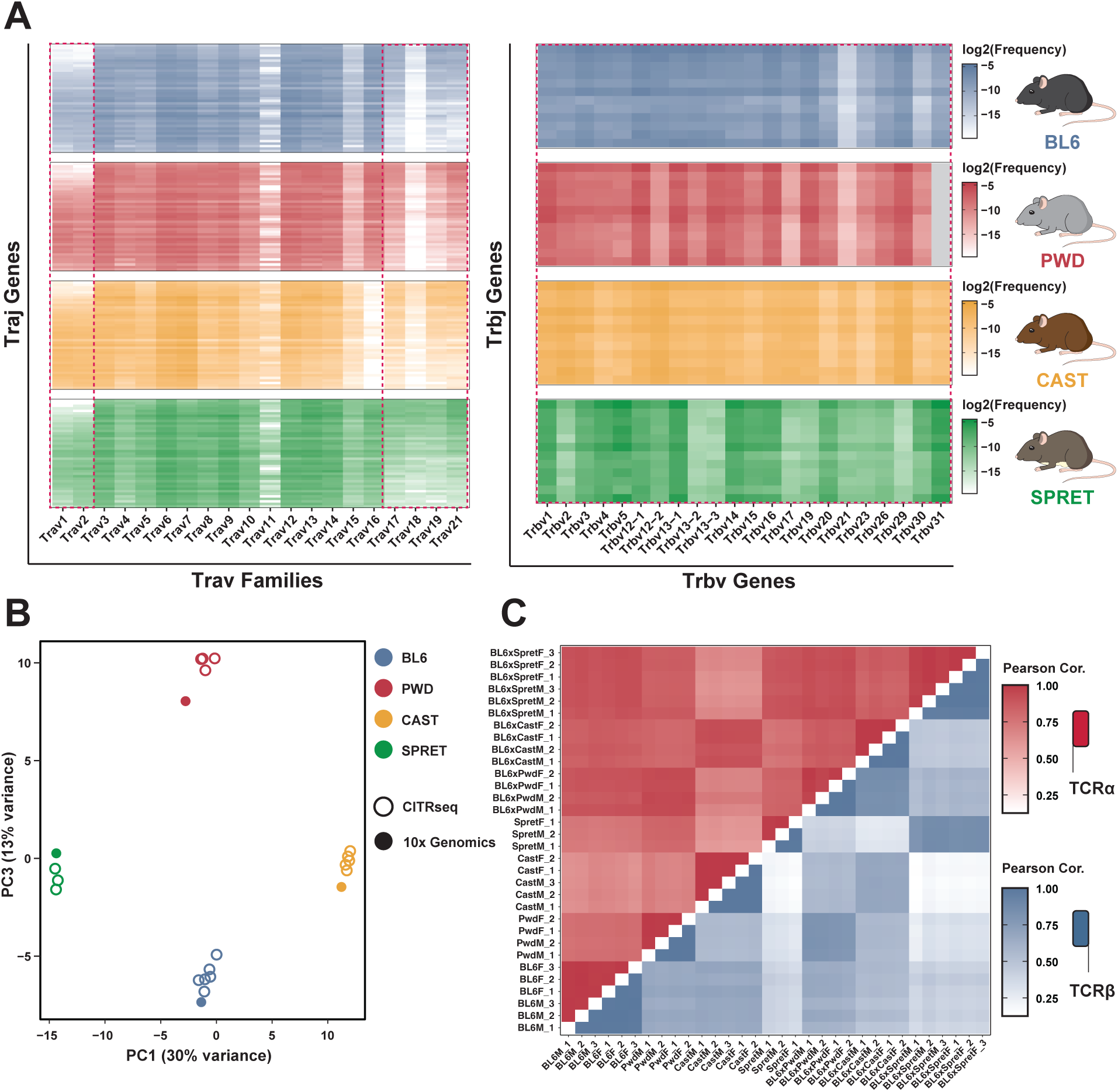
Species-specific V- and J-gene usage patterns in different mouse species. **(A)** V-J usage frequency heatmaps. Heatmap shows the frequency (log_2_) of Vα-family + Jα-gene (left) and Vβ-gene + Jβ gene (right) usage of all functional TCRs in T cells across the four different mouse species (intra-species mean). Red boxes contain V-genes with one-to-one orthology in all four mouse species. J-genes are displayed in the order of their location within the locus (3’ to 5’; see methods for full list). In PWD *Trbv31* is excluded due to failure of amplification during multiplex PCR. **(B)** Principal component analysis (PCA) of combined Vα-Jα and Vβ-Jβ usage across all four mouse species in different samples generated using CITR-seq (empty circles) or 10x Genomics Single Cell Immune Profiling (filled circles). Samples generated using both methods cluster by genotype. **(C)** Pearson correlation of inter-individual V-J gene usage in TCRα and TCRβ chains in all 32 individuals analyzed using CITR-seq in this study.

Further to the previous validation effort, we have also generated matching V(D)J usage profiles using the commercial 10x Genomics platform. Similar to BL6 mice, we observed excellent correlation of V-J gene usage frequencies in both approaches (Pearson correlation: PWD r = 0.89, CAST r = 0.88, SPRET r = 0.92) and only identified two V-gene segments that were not recovered in the CITR-seq dataset compared to the 10x Genomics dataset (*Trbv-31* in PWD, and *Trbv-24* in CAST/SPRET). The main difference between the platforms is that with CITR-seq we recovered on average ∼160,000 T cells carrying a productive and paired TCR per experiment vs. 2,300 using the 10x Genomics platform.

Next, we performed principal component analysis (PCA) on combined TCRα and TCRβ V-J pairings across all CITR-seq replicates from each species alongside the 4 samples generated using 10x Genomics Immune Profiling, subsampling each sample to 5,000 TCRα and TCRβ chains each due to the lower throughput of the latter (**see methods**). Overall, samples are clustered strongly by species (PC1-PC3; **Fig. 3B**, **Fig S3E**) with only 6% of cross-sample variance explained by the technique (PC4, **Fig. S3E**). Mean intra-species V-J usage was highly correlated across samples for both TCRα and TCRβ (Pearson: r = 0.987 +/- 0.052 stdev vs. r = 0.991 +/- 0.044 stdev) (**Fig. 3D**). Across samples the average V-J segment usages were more correlated for TCRα chains (r = 0.83 +/- 0.119 stdev) than TCRβ chains (r = 0.59 +/- 0.244 stdev) which is consistent with the difference in overall diversity across both chains described earlier (**Fig. 2E**). Therefore, we conclude that in the four different mouse species, V-J usage showed distinct genotype specific patterns primarily in Vβ-genes.

### Thymic selection shapes TCR repertoire V-segment usage

Thymic selection ensures that T cells expressing TCRs with either too weak (positive selection) or to strong (negative selection) self-MHC binding properties fail to progress in the maturation process and are thus depleted from the repertoire. In effect, by collecting TCRs from peripheral (e.g., spleen-derived) T cell populations for CITR-seq, we report here the mature TCR repertoire after thymic selection. Here, we also emphasize a distinction between functional vs. non-functional TCR chains. This is because during V(D)J-recombination, random insertions and deletions of nucleotides at gene-segment junctions, often introduce frameshifts or premature stop-codons. These result in transcripts representing non-functional TCRs. However, mature T cells with an in-frame (IF) TCR often still retain active transcription of an out-of-frame (OOF) TCR from its second allele that is ultimately degraded, e.g., via non-sense mediated decay [52, 53]. This presented us with an opportunity to estimate the generative usage probability of gene segments, independent from the effects of positive or negative thymic selection (see also [54]). Crucially, our use of an inbred panel of species should result in an unchanged, homozygous MHC background resulting in a consistent thymic selection regime.

Across all 32 CITR-seq sample we found 4.58×10^6^ (24.4% of total transcripts) transcripts that contain frameshifts or premature stop-codons with an average per transcript UMI count of 1.78 (compared to 4.87, two-sample t-test, *P* < 0.001). To evaluate the effect of thymic selection on TCR repertoires across the different species, we compared V- and J-gene usage in OOF (pre-selection) and IF (post-selection) TCRs (**Fig. 4A**). We observed that most V(D)J genes show similar frequencies in pre- and post-selection repertoires, summarized by normalized Shannon diversity index (nSDI, **see methods**), a measure of entropy (Vα (BL6 and PWD) as well as Jα (no significant changes) and Jβ (PWD and SPRET; **Fig. S4B**)). Again, the strong exceptions reside mostly within Vβ-genes: we observed significant differences in Vβ gene usage frequencies in all four species (**Fig. 4B**, paired t-test *P* < 0.05). The strongest absolute reduction of nSDI was observed in SPRET (-0.15) and PWD (-0.06), indicating significantly biased Vβ-segment usage in post-selection repertoires. For specific Vβ-segments, we found striking differences between pre- and post-selection repertoires, e.g., an average ∼60-fold reduction in *Trbv13-2* usage frequency in SPRET post-selection repertoires (**Fig. 4C**). Notably, these extreme fold changes were mostly present in Vβ genes that showed strong cross-species frequency difference (e.g. *Trbv-2*, *Trbv12-2*, *Trbv13-2*, *Trbv17*, *Trbv21*) as shown before (**Fig. 3A**). We interpret the striking reduction in usage for these Vβ segments to be strongly suggestive of segment rejection during thymic selection.

**Figure 4:**
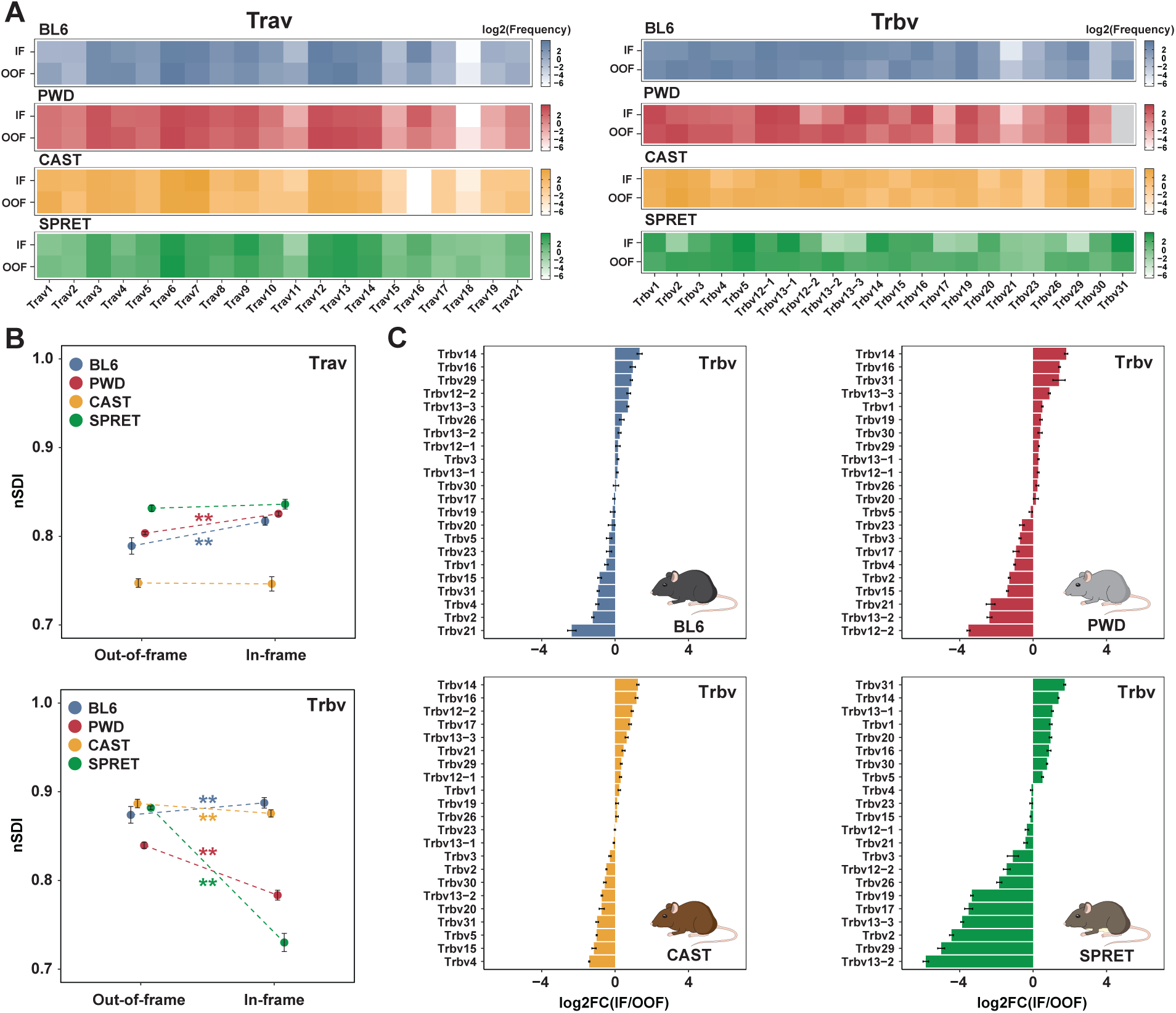
Thymic selection shapes V-gene usage. **(A)** Vα family (left) and Vβ gene (right) usage frequency (log_2_) heatmaps. Heatmaps show the mean intra-species V-usage in in-frame (IF) and out-of-frame (OOF) TCRs across all T cells. **(B)** Mean intra-species entropy in Vα-usage (top) and Vβ-usage (bottom) distributions calculated using the normalized Shannon diversity index (nSDI) for OOF and IF TCRs (error bars indicate the standard deviation in species replicates, significance calculated using a paired t-test, * *P-value* < 0.05). **(C)** Log_2_ fold-changes (log2FC) in Vβ gene usage frequencies between IF and OOF TCRs across the different mouse species (error bars indicate the standard deviation in species replicates).

While the most extreme differences in Vβ-segment usage tend to be species specific, we also observed common trends shared across all four species. For instance, we observed *Trbv-2* frequencies to be consistently lower in post-selection than pre-selection repertoires across all species (log2 FC IF/OOF; BL6: -1.2, PWD -1.3, CAST -0.5, SPRET -4.4). While thymic selection acts only to remove T cells from maturation, such that the absolute number of TCRs containing a particular gene segment only decrease from pre- to post-selection repertoires, in relative terms, a given segment can be overrepresented in the final, mature repertoire through thymic selection. One such example was *Trbv-14* whose relative contribution to the TCR repertoire was higher in all post-selection repertoires across species (log2 FC IF/OOF; BL6: 1.3, PWD 1.8, CAST 1.24, SPRET 1.38).

In summary, we show that thymic selection exerts an effect on the composition of the TCR repertoire by distorting usage frequencies in all segments across all four species, but its effect is most notable in Vβ-genes, in particular in the reduction of particular Vβ-genes (e.g. *Trbv13-2*, *Trbv2*, *Trbv12-2* etc.) in PWD and SPRET, likely due to strong rejection during positive thymic selection.

### Allele-specific V/J segment usage in F1 Hybrids revealed by patterns of thymic selection

In contrast to outbred individuals, assaying V(D)J gene usage in inbred mouse strains benefits from consistent thymic selection, thanks to the homozygous MHC-allele background. If so, the observed OOF-IF profile should shift in individuals carrying alternative MHC-haplotypes (see also [55, 56]). This raises further the tantalizing possibility that, depending on the actual MHC-haplotype, there may be different outcomes associated with positive vs. negative thymic selection. To test our hypothesis, we generated F1 hybrids from crosses of BL6 with each of the three other mouse species (BL6xPWD, BL6xCAST, BL6xSPRET). This gave us a powerful tool to track how the two otherwise distinct sets of species-specific V(D)J gene repertoires may be shaped by thymic selection in the respective heterozygous MHC allele state.

We first compared V(D)J usage frequencies in F1 hybrids with the respective frequencies in the parental species (**Fig. 5A** for V-genes and **Fig. S5A** for J-genes). Similar to our previous analysis, we see the most differences across Vβ-genes, in both directions: Vβ- genes can be significantly more abundant (e.g., *Trbv1*) or less abundant in F1 hybrids than in either parent (e.g., *Trbv12-1* and *Trbv12-2*; Wald-test; *P* < 0.01; **Fig. S6A-D**). To analyze the general trends across Vα, Jα, Vβ and Jβ frequencies between parental lines and their F1 hybrids, we classified their relative V(D)J gene usage frequencies into five broad categories: conserved, additive, dominant, over- and under-dominant (**Fig. 5B, see methods**). In Vα and Jα genes, 78.6% of genes only show modest frequency changes (<1%) relative to both parents. Over- and underdominance (>1% higher/lower frequency than both parents, respectively) are only seen in genes in the TCRβ chain and mostly in Vβ-genes. Notably, we observe overlaps in the identity of over-dominant (e.g., *Trbv1*) and under-dominant (e.g., *Trbv12-1*) Vβ-genes across all three hybrids. Collectively, V(D)J gene frequency changes between F1 hybrids and the parental lines are predominantly observed in the TCRβ chain.

**Figure 5:**
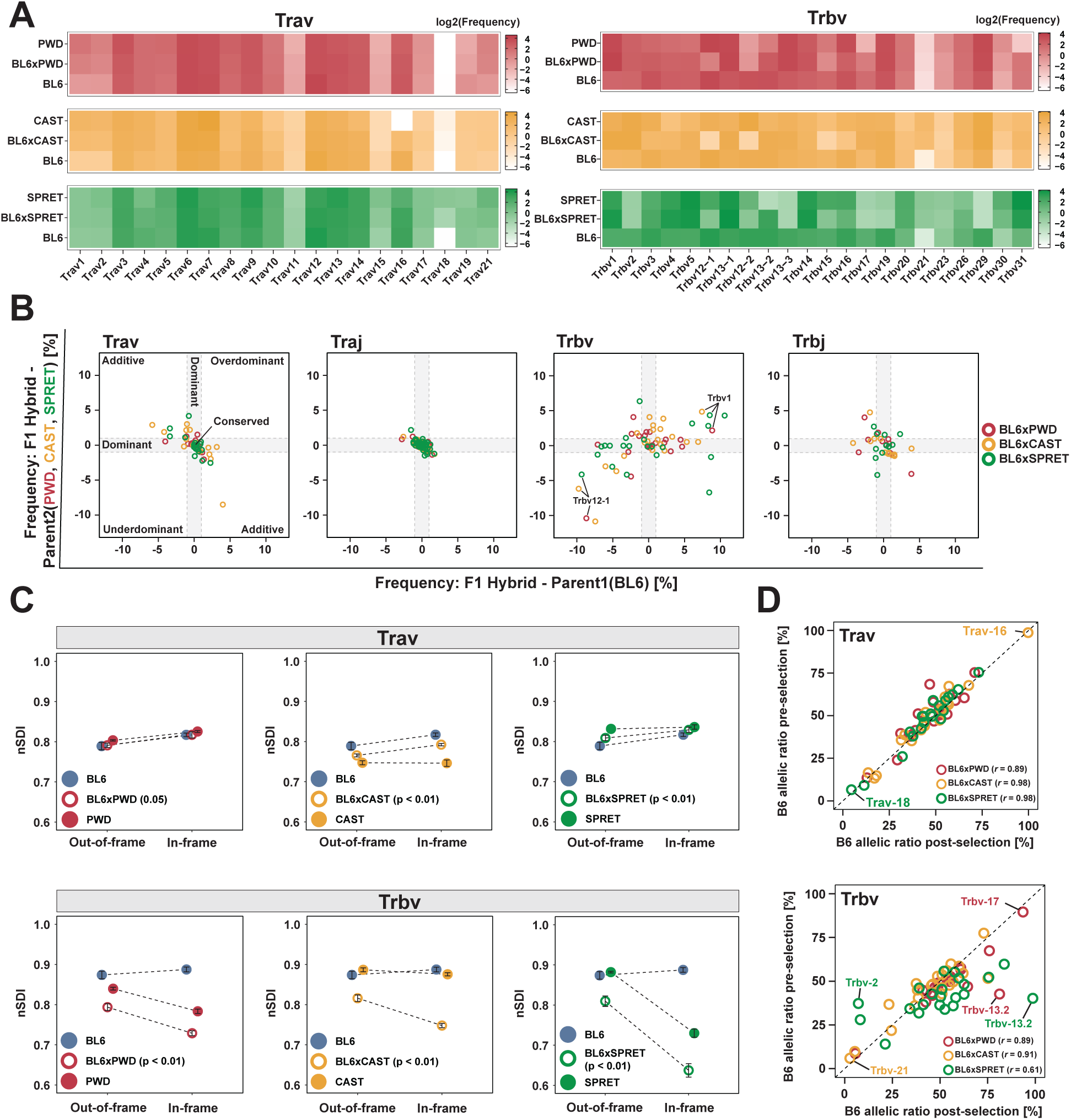
V-gene usage is more restricted in F1 hybrids and shows allele specific usage biases. **(A)** Vα family (left) and Vβ gene (right) usage frequency (log_2_) heatmaps of in-frame TCRs in F1 hybrids and their respective parental species. **(B)** Relative frequency changes of V(D)J gene usage in F1 hybrids and the respective parental species (x-axis: F1 hybrid – BL6 and y-axis: F1 hybrid – PWD, CAST or SPRET) categorized into mode of inheritance. Conserved (center), dominant (grey area), additive (top left and bottom right quadrant), under-dominant (bottom left quadrant) and over-dominant (top right quadrant; see methods). Each circle represents a Vα-family, Jα-gene, Vβ-gene or Jβ-gene. **(C)** Comparison of entropy of V-usage distribution in F1 hybrids and the respective parental species calculated using the normalized Shannon diversity index (nSDI) for OOF (left) and IF (right) TCRs (error bars indicate the standard deviation in species replicates, significance tested for F1 hybrid IF vs OOF contrast using paired t-tests, *P-value* < 0.05). **(D)** Analysis of biased V gene allele usage in F1 hybrids. Plots show the percentage of BL6 Vα family alleles and Vβ gene alleles in post- (x-axis) and pre-selection (y-axis) TCRs. Each circle represents a Vα-family (top) or Vβ-gene (bottom). Pearson-correlation was calculated for post- and pre- selection V gene usage.

We then calculated the nSDI for pre- and post-selection repertoires in the F1 hybrids and compared them to the previously calculated nSDI values in the parental species (Trav and Trbv **Fig. 5C** Traj and Trbj **Fig. S5B**). Across all F1 hybrids, we see significantly reduced nSDI values for Vβ-gene frequencies (*P* < 0.05; paired t-test). The increase in unevenness between pre- and post-selection repertoires are consistently greater in F1 hybrids compared to their parental species, suggesting that thymic selection introduces stronger biases on Vβ-gene usage in F1 hybrids relative to their respective parental species. Interestingly, we observed a constant increase of nSDI values in post-selection compared to pre-selection Vα gene frequencies in F1 hybrids (*P* < 0.05 in BL6xCAST and BL6xSPRET).

Next, we took advantage of our ability to assign V(D)J genes in F1 hybrids in an allele-specific manner to identify potential biases towards usage of one parental allele. We compared the allelic ratios of V(D)J genes in pre- and post-selection repertoires (V genes: **Fig 5D, Fig S5D** and J genes: **Fig S5C**). We found significant allelic biases in Vβ-genes in post-selection repertoires that were not observed in the pre-selection repertoire. For example, in pre-selection repertoires of BL6xSPRET hybrids, ∼60% of *Trbv13-2* usage was assigned to the SPRET allele, whereas in post-selection repertoires this rate dropped to ∼1%. Therefore, while in BL6xSPRET hybrids the *Trbv13-2* allele was frequently recombined during V(D)J recombination, it was almost completely rejected during thymic selection. The almost exclusive selection of one parental allele in a heterozygous MHC haplotype and a common *trans*-environment, provides strong evidence that this selection process is primarily determined by genetically encoded polymorphisms in the underlying Vβ gene. Similarly, we saw that while Traj35 pre-selection frequencies are balanced between parental alleles (Percent of BL6 alleles: PWD 49%, CAST 44%, SPRET 50%), the BL6 allele was substantially less frequent in post-selection repertoires (Percent of BL6 alleles: PWD 25%, CAST 26%, SPRET 25%; **Fig S5C**).

Apart from these strong exceptions, allelic bias is strongly correlated in pre- and post-selection repertoires for most V(D)J genes (see Pearson correlation in **Fig. 5D** and **S5C**). Genes that show strong frequency differences between both parental species (*Trav16* and *Trbv21* in BL6/CAST, *Trav18* in BL6/SPRET or *Trbv17* in BL6/PWD) often show strong F1 allelic bias towards usage of the respective parental allele that had a higher frequency in the pure contrast (**Fig. 5D**). We therefore conclude that the (generative) pre-selection biases observed between species are primarily controlled by linked factors acting in *cis*, e.g., polymorphisms in the RSS sequences that influence the recombination likelihood of a particular gene during V(D)J recombination.

### Composition and diversity of the paired CDR3αβ repertoire depends on an individual’s genotype

Next, we addressed CDR3 diversity 32 samples, representing seven different genotypes. We reasoned that given the cumulative bias in gene segment usage, availability, as well as genetic differences, we should observe distinct CDR3 amino acid motif repertoires. To test this hypothesis, we first compared the CDR3α, CDR3β and CDR3αβ diversity in a set of 100,000 T cells sampled randomly from each individual (**Fig. 6A**). We found that, across all comparisons, F1 hybrids show an increased number of unique CDR3 sequences relative to their respective parental species. In single CDR3α motifs, we see significant differences in diversity within the parental species, which is in line with the observed differences in locus structure of the TCRα loci across these mice (e.g., ∼50 fewer Vα segments in CAST compared to all other species; pairwise t-test; ** = *P* < 0.01; * = *P* < 0.05). These parental diversity differences are not recapitulated in the F1 hybrids. Instead, the absolute diversity increases with the increasing evolutionary divergence of the parental species.

**Figure 6:**
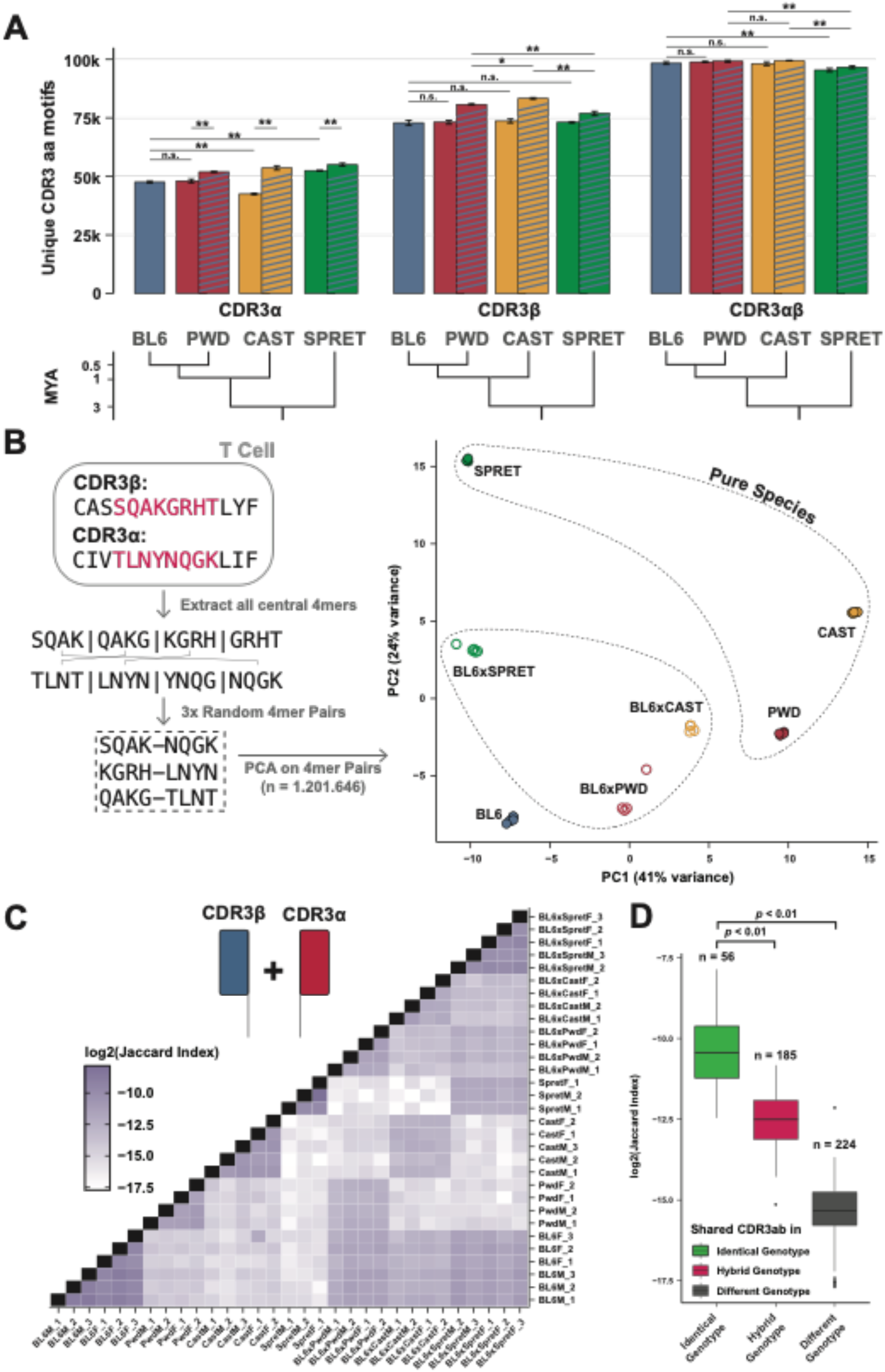
CDR3 motif diversity and sharing depends on an individual’s genotype. **(A)** Mean count of unique CDR3α (left) CDR3β (middle) and paired CDR3αβ (right) amino acid motifs in a set of 100,000 randomly sampled TCRs from each individual grouped by genotype (error bars indicate the standard deviation in species replicates, significance calculated by pairwise t-tests with * *P-value* < 0.05 and ** *P-value* <0.01). Single-color bars represent the parental species, diagonally striped bars represent the respective F1 hybrids. The phylogenetic tree (bottom) shows the evolutionary divergence of parental species. **(B)** Analysis of paired 4mers extracted CDR3αβ motifs. For each paired CDR3αβ amino acids sequence all possible 4mers were extracted and subsequently, 3 random 4mer pairs (one from the CDR3α and one from the CDR3β sequence) were generated (left panel). The combined filtered (see methods) count matrix of 4mer pairs from all 32 individuals was then used for PCA analysis. Samples cluster based on the sample genotype. **(C)** Overlap of paired CDR3αβ amino acid motifs between all 32 CITR-seq samples calculated using the Jaccard index (log_2_; see methods) **(D)** Based on overlap of genotypes all samples were grouped into identical genotype (within species, e.g., BL6F_1 and BL6M_1), hybrid genotype (50 % identical genotype, e.g., CAST and BL6xCAST) and different genotype (completely unrelated individuals, e.g. PWD and SPRET). Boxplot shows the calculated Jaccard index values (log_2_) in each respective group (significance tested using Wilcoxon rank-sum test, *P-value* < 0.01).

A strikingly different picture emerged for the CDR3β motifs. Importantly, V(D)J segments in the TCRβ locus follow a strict one-to-one orthology across all parental species. Accordingly, single-chain CDR3β diversity showed little variation across the parental species. In contrast, F1 hybrids showed much greater variation and generally display greater TCR diversity than observed in the repertoire of either parent (pairwise t-test; **= *P* < 0.01; * = *P* < 0.05). A possible explanation for this is the strictly restricted selection of Vβ segments during thymic selection in the TCRβ chain. We observe remarkable diversity of paired CDR3αβ across all genotypes, with an average of 98.2% of all motifs being observed only once in each set of 100,000 motifs. Notably, among F1 hybrids, the lowest diversity is observed in BL6xSpret hybrids despite the highest evolutionary divergence in the respective parental species.

Due to random insertions and deletions at the segment junction sides, the highest amino acid diversity within CDR3 motifs is observed in the central region of the peptide chains (**Fig**. **S7A**). It has been shown that this central region overlaps the region of highest antigen proximity in TCR-MHC complexes (position 107-115 according to IMGT nomenclature [57]) and therefore contributes most to the antigen specificity of the underlying TCR [6]. Further, the same study also provided evidence that antigen-specificity is defined by specificity-groups of similar amino acid motifs within TCRs. With this in mind, we analyzed germline-encoded differences in the central motifs across all mice. Because the same antigen might be recognized by several similar TCRs rather than just one CDR3αβ motif we first generated amino acid 4mers from both CDR3 motifs of TCRs of individual cells (**Fig 6B**). We then identified a list of 1,201,646 common 4mers across all genotypes (**see methods**). Next, we performed PCA analysis based on the abundance of all 4mer pairs across all 32 individuals. We see that 4mer pairs are strictly clustered according to the underlying genotype of each sample (**Fig. 6B**). This pattern was also observed in the corresponding analysis on single-chain derived unpaired 4mers (**Fig S7B**).

Comparison of TCR repertoires often involves the analysis of shared “public” CDR3 motifs. Typically, this type of analysis addresses motif sharing within single chains across repertoires. While these comparisons might provide information on the generative probability of distinct single-chain CDR3 motifs across individuals, the missing CDR3 motif in the second chain makes it challenging to identify potential shared TCR responses to antigens. Here, we utilized CITR-seq’s large set of more than 5 million paired CDR3 motifs to analyze motif sharing across all individuals. In total, we identified 25,894 (∼0.5% of all motifs) paired motifs with identical amino acid sequence, observed in different individuals. Across single chains, sharing of identical amino acid sequences was more common with 264,088 shared CDR3α (∼36.7% of all unique motifs) and 469,827 shared CDR3β (∼ 27.2% of all unique motifs) motifs observed in at least two individuals. Notably, we found 1696 CDR3α and 644 CDR3β amino acid motifs that were observed in all 32 individuals, while identical CDR3αβ pairs were at most observed in 12 individuals (**Fig. S7C**)

To test whether the extent of paired CDR3αβ motif sharing was higher than expected by chance, we shuffled the α- and β-chains within each individual and then re-calculated the count of shared motifs. We saw that the observed sharing count is about 4-fold higher than the mean across 100 permutations of αβ-chains shuffled samples (mean: 6182 shared CDR3αβ motifs across shuffled pairs, permutation test *P-value* < 0.01). Next, we analyzed whether the extent of sharing in CDR3αβ motifs is dependent on the underlying genotype of each sample. To account for the variance in sample size across all samples, we calculated the Jaccard Index of repertoire sharing using paired CDR3αβ motifs (**Fig. 6C, see methods**). We observed that motif sharing is significantly higher across samples of identical genotypes (56.7% of all shared motifs), compared to individuals with partially shared genotypes (F1 hybrid samples, 32.0% of all shared motifs) and especially in contrast to completely unrelated individuals (11.3% of all shared motifs; Wilcoxon rank sum test *p-value* < 0.01) (**Fig. 6D**). While we caution here that our use of inbred individuals may differ from the usual comparison contexts with CDR3 motifs, nevertheless, the extent of sharing across fully unrelated individuals led us to conclude that an individual’s genotype contributes significantly to the final TCR repertoire. Additionally, public TCR responses are far more likely to be observed across related individuals than unrelated individuals.

## Discussion

Production and maintenance of large and diverse repertoire of TCRs is crucial for a functioning adaptive immune system. For decades researchers have now accumulated insights into the generative process, the size and overlap, as well as associations to disease states of TCR repertoires. High-throughput sequencing technologies have reached sufficient sensitivity and throughput to capture reasonable portions of an individual’s TCR repertoire. Yet they still suffer from severe limitations in the face of the extreme diversity of TCR repertoires. To date, arguably the most limiting of these factors is the requirement for single-cell resolution to link both TCR chains of the heterodimeric αβ receptor to the T cell of origin. With few (mostly non-commercial) exceptions, single-cell TCR sequencing methods suffer from low-throughput (10^3^-10^5^ T cells) and high cost (reviewed here [58]). Pit against the vast TCR repertoire diversity, especially in naïve repertoires, those technologies often capture only a tiny fraction of an individual’s repertoire.

In this study we present CITR-seq, a high-throughput low-cost single-cell TCR sequencing method that overcomes many of these limitations. We use CITR-seq to generate TCR repertoires of four evolutionary divergent inbred mouse species and t their respective F1 hybrids, covering more than 9 million T cells with 76% successful αβ-pairing rate.

We first identified large differences in V(D)J gene usage across the different mouse species, with very high within-species consistency for both TCR chains. While the arrangement and number of genes in the TCRβ locus are conserved across all species, the TCRα locus has undergone complex rearrangements leading to triplications and inversions of Vα gene clusters. As a result, the number as well as their relative distance to Jα genes varies substantially between Vα genes of the different mouse species. We observed that at the TCRα locus in CAST, in which the Vα locus was contracted by 0.6 Mb, the distal Vα genes showed significantly higher segment usage compared to the other species. Considering the progressive 3’ to 5’ recombination of V-J segments [59], we interpret this as evidence for a direct relation of gene segment locus size and chromosomal position dependent usage frequency.

Due to the very conserved arrangement of TCRβ genes, the tightly enforced allelic exclusion as well as prevention of continuous rearrangements, the relative position of genes should contribute less to biases in the TCRβ gene usage across species. Nevertheless, we see that the relative fold-changes in gene usage of Vβ genes can be extreme, with up to 60-fold difference between different mouse species.

We show that many of those extreme gene usage differences are introduced during thymic selection by comparing pre- vs post-selection repertoires. We use the nSDI of segment usage to demonstrate that thymic selection primarily acts on Vβ segments and show that their generative frequency immediately after V(D)J recombination is more similar across different mouse species than the actual usage frequencies observed in mature and selected TCR repertoires. Critically, we observed that many of the Vβ genes that are rejected during thymic selection, contribute identical amino acids to CDR3 motifs compared to other Vβ genes that do not significantly change in frequency in pre- vs. post-selection repertoires. Thus, we hypothesize, that the rejection of those Vβ genes is unlikely to be enforced during negative selection as a consequence of strong affinity to a self-MHC complex. Rather, it is reflective of their particular germline-encoded ability to bind MHCs evaluated during positive selection. This hypothesis is further supported by the fact that we did not observe categorical rejection of J-segments, that are though to mostly contribute to the antigen specificity of a TCR rather than its ability to bind to MHCs. Further experiments, where both Vβ and MHC components can be experimentally controlled, may be able to shed light on the mechanism underlying our observation.

We also used F1 hybrids of inbred mouse species, as a powerful tool to evaluate the thymic selection of TCRs in a defined heterozygous MHC haplotype. In those hybrids, two sets of V(D)J genes are exposed to a common *trans*-environment that subject both to a common pre- and post-selection repertoires regime. As a general trend, we see that most V(D)J genes show conserved usage frequencies relative to the parental species, or alternatively, in the case of substantial differences between the parental species, exhibit intermediate (*additive*) gene usage frequencies. These general patterns are far less pronounced for Vβ gene usage frequencies of F1 hybrids. We see that the selection against particular Vβ genes mostly resembles the patterns seen in the parents, with additional rejections of particular genes that were frequent in both parents. By utilizing our species-specific V(D)J references, we were able to disentangle the usage frequencies of particular alleles in F1 hybrids. We provide examples of Vβ-genes with balanced allelic ratios in pre-selection repertoires and striking allelic biases in post-selection repertoires. The nearly mono-allelic usage of particular Vβ-genes as a consequence of thymic selection in a defined heterozygous MHC allele state in F1 hybrids, provides strong evidence that the rejection of particular Vβ-gene alleles is based on genetically encoded polymorphisms. To the best of our knowledge, such extreme cases of allele-specific Vβ-genes selection have not been described before. This finding has important implications for the ongoing debate about whether binding to MHCs is an inherent and germline-encoded feature of TCRs that progressively co-evolves, or alternatively, MHC restriction of TCRs is enforced by TCR co-receptor signaling involved in TCR-MHC complex formation. Due to the common *trans-*environment during thymic selection of TCRs, the strong allelic biases of particular Vβ genes can hardly be explained by co-receptor signaling and thus should reflect the inherent ability of particular Vβ gene alleles to bind MHCs originated from heterozygous alleles in the F1 hybrids. Consequently, we hypothesize that TCR-MHC binding is a co-evolutionary process mediated by changes in amino acid sequences of V gene regions and MHC alleles that facilitate complex formation. In this context, the highly variable germline-encoded CDR1 and CDR2 regions of TCR V-genes have been shown to be crucial for altering TCR-MHC binding strength.

A secondary consequence of the co-evolution of TCR-MHC binding would likely be the increased rate of TCRs that exhibit insufficient or overly strong affinity to MHCs in hybrids between highly divergent parents. Indeed, as shown in this study, thymic selection had the strongest effect on Vβ genes in BL6xSPRET individuals in which the respective parental individuals had the highest degree of evolutionary divergence.

We further show that biases that are consistent in pre- and post-selection repertoires mostly reflect selection independent gene usage frequency differences observed in the parents. For instance, TCR consisting of *Trbv21* are extremely rare in BL6 (0.03% of TCRs) but much more frequent in CAST (3.0% TCRs) with minor differences in pre- and post-selection frequency. In BL6xCAST F1 hybrids, 2.2% of all TCRs consist of *Trbv21* with an allelic ratio of 97.3% of CAST alleles and only 2.7% BL6 alleles. Therefore, gene segment frequency biases are mediated through *cis*-effects in the absence of any additional biases introduced by thymic selection. For instance, polymorphisms in the RSS in between V(D)J genes could bias the recombination efficiency of particular gene segments.

What are the consequences of the observed frequency and selection biases across the different mouse species for the total diversity within the TCR repertoires? To answer this question, we evaluated CDR3 motif diversity in single chains as well as paired TCRs. CDR3α diversity varies most across the pure species, which is likely caused by the severe rearrangements and consequently different number of functional gene segments in the Vα cluster. We generally observe minor frequency changes of Vα families in pre- and post-selection repertoires, indicating that those gene segments are subject to less stringent thymic selection. As a consequence, F1 hybrids can make full use of both parental sets of V(D)J genes, which likely leads to the correlation of increased CDR3α diversity with increased evolutionary divergence of parental species. Here, we note that grouping of Vα genes by their respective families might mask the rejection of particular genes during thymic selection. While this potentially impacts the gene usage frequency differences across the species, it does not bias the comparison of total CDR3α diversity. The single-chain CDR3β motif diversity is extremely similar across pure species, which is in line with the one-to-one orthology of V(D)J genes in the TCRβ locus. In contrast to this, we see substantial differences in CDR3β diversity in the hybrids. Based on the observed impact of thymic selection on Vβ genes, we hypothesize that the observed differences in CDR3β diversity in F1 hybrids result from a trade-off between the diversity of parental V(D)J gene sets and the increased likelihood of gene segment rejection during thymic selection, which should correlate with the increasing evolutionary divergence of parental species. While in this study, thymic selection is evaluated in a fixed and genotype-specific MHC-haplotype set-up, it has been shown that increased intra-individual MHC diversity is associated with increased rates of T cell depletion during thymic selection [60, 61]. Given that HLA allele frequencies vary substantially across human populations [62], we assume that the general trends observed in this study would therefore also apply in the context of CDR3 diversity evaluation in evolutionary divergent outbreed populations exhibiting diverse MHC haplotypes.

To the best of our knowledge, the present study analyzes the largest set of paired αβ-TCR to date. Especially in the context of TCR repertoire analysis of antigen inexperienced naïve T cells we benefit greatly from the scale of our dataset. Sharing of identical CDR3αβ motifs is rare but about 4-fold higher than expect by chance. Additionally, shared motifs are found at significantly higher rates in related individuals compared to unrelated individuals. Importantly, the increased sharing rate of paired CDR3αβ motifs analyzed in this study is not limited to the comparison of 100% identical motifs. Because similar CDR3αβ motifs might recognize identical antigens and similar antigens might be recognized by a range of similar CDR3αβ motifs we used a kmer-based approach to emphasize the similarity of paired CDR3αβ motifs in species of identical genotypes. We showed that 4mers originated from the central region of paired CDR3αβ motifs exhibit remarkably similar frequencies in species with identical genotypes relative to unrelated individuals. We therefore conclude that the combined effects of differences in TCR locus structure, V(D)J recombination frequencies and biases introduced by thymic selection, collectively shape the TCR repertoire in a genotype-specific manner.

This also has important implications for our understanding of public TCR motifs with potential disease associations. The number of shared CDR3 motifs in individuals with diverse MHC haplotypes is representative of those TCRs, that are selected by the specific set of MHCs in the sampled individuals. Public CDR3 motifs should therefore always be cataloged in the specific MHC haplotype context they have been observed in to allow for the comparison of such public motifs across different studies. Additionally, public TCR responses are often evaluated in common disease context, such as cytomegalovirus (CMV) and Epstein-Barr virus (EBV) [63-65]. Since large parts of human populations are persistently infected by those pathogens, a broad range of MHC haplotypes should have evolved to effectively present EBV- and CMV-derived peptides. Consequently, EBV- and CMV-associated CDR3 motifs might be more public compared to CDR3 motifs that specifically recognize less frequent pathogenic peptides.

Immune receptor diversity is one of the most characteristic and important features of adaptive immunity. While the generation of diversity is in large parts driven by stochastic events, the present study highlights important genetic contributions to TCR diversity. We show that the number of functional V(D)J segments, their *cis*-regulated recombination frequency as well as MHC haplotype dependent thymic selection, collectively generates TCR repertoires that are significantly more similar within than across genotypes.

## Methods

### Mice

All mice were housed in the animal facility of the Friedrich-Miescher Laboratory of the Max-Planck Society. Experiments were performed under license issued by the local competent authority (EB 01/21 M). Spleens were collected from mice aged 9-11 weeks. The following mouse strains were used in the experiments: C57BL/6J (The Jackson Laboratory, Strain #: 000664), CAST/EiJ (The Jackson Laboratory, Strain #: 000928), SPRET/EiJ (The Jackson Laboratory, Strain #: 001146), PWD/PhJ (The Jackson Laboratory, Strain #: 004660) as well as their respective F1 hybrids (C57BL/6J x SPRET/EiJ/CAST/EiJ/ PWD/PhJ). Male and female mice of all strains were used.

### Isolation of CD8a^+^ T-cells

Spleens of euthanized mice were collected and placed on a 40µm cell-strainer. Spleens were then pressed through the strainer using the backside of a syringe plunger. After thorough rising of the cell-strainer using ice-cold PBS, the flow-through was centrifuged at 400xg 4°C for 10 minutes in a swing-bucket centrifuge. Afterwards, supernatant was carefully discarded, and the cell pellet was resuspended in 1ml ice-cold PBS + 2% FBS. Isolation of CD8a^+^ T-cells was then done using the “Dynabeads™ FlowComp™ Mouse CD8 Kit” (Invitrogen, 11462D) according to the manufacturer’s instructions. Pre-enriched cells were then stained using anti-CD4 BV510 (Bio Legend, 100553) and anti-CD8 PerCP-Cy5.5 (Bio Legend, 155013) in 500µl PBS + 2% FBS for 15 minutes on ice. Afterwards, cells were centrifuged at 400xg 4°C for 5 minutes. Supernatant was discarded and cell pellet was resuspended in 500µl ice-cold PBS + 2% FBS. This washing step was repeated once before final resuspension in 1 ml ice-cold PBS + 2% FBS. Cells were then further purified by fluorescence activated cell sorting (Fig. S1A). Depending on the size of the spleen (approx. 20mg in SPRET and up to 100mg in BL6) between 1×10^6^ and 5×10^6^ CD8+ T-cells were isolated from each spleen. Isolated T-cells were immediately transferred to prepared tissue culture dishes or used as primary cells for CITR-seq experiments.

### Tissue Culture

Tissue culture of isolated CD8^+^ T-cells was done as described by Lewis et al. [66]. Briefly, 6-well plates were coated with 0.5µg/ml anti-CD3 and 5µg/ml anti-CD28 in 3ml PBS at 4°C overnight. Before seeding the isolated CD8^+^ T-cells, plates were washed twice with PBS. Cells were cultured in RPMI 1640 medium (ThermoFisher, 11875093) supplemented with 10% FBS, 1% GlutaMAX (ThermoFisher, 35050061), 1% penicillin/streptomycin (ThermoFisher, 15140122), 0.1% 2-mercaptoethanol (ThermoFisher, 21985023) and 0.1% human recombinant insulin (ThermoFisher, 12585014) at 37°C, 5% CO_2_. After 20 hours cells were washed once with culture medium and then carefully detached from plate by repeatedly flushing the plates with a P1000 pipette. The cell suspension was then centrifuged at 400xg, RT for 5 minutes. Afterwards cell pellet was resuspended in 1ml PBS.

### CITR-seq protocol

#### Oligonucleotides for barcoding

Two rounds of barcoding, each with 192 unique DNA barcodes are performed in CITR-seq. To prepare the barcoding plates in each well of two 96-well plates one unique round 1 top-stand oligo and one corresponding round 1 bottom-strand oligo were diluted in 10µl annealing buffer (10mM Tris pH 8, 50mM NaCl and 1mM EDTA). Top-strand round 1 oligos are partially complementary to the 5’ overhang of the RT primers and anneal to the complementary sequence of the round 1 bottom-strand including the 7bp barcode sequence. Round 1 bottom-strand oligos contain a common 3bp 5’ phosphorylated linker overhangs (“TCT”). The same procedure was repeated for two 96-well round 2 barcoding plates. Round 2 top-strand oligos contain a 3’-linker sequence (“AGA”) complementary to the 5’ linker sequence of round 1 oligos. Further, it contains another unique 7bp DNA barcode and the standard Illumina TrueSeq i7 sequencing adapter (Illumina, see document: 1000000002694). Round 2 bottom-strand oligo is complementary to its respective round 2 top-strand mate but lacks the 3bp linker sequence.

Oligos are used at the following concentrations: For each well of round 1 plates: µM of round 1 bottom-strand and µM of round 1 top-strand. For round 2 plates: µM of round 2 bottom-strand and µM of round 2 top-strand. Prior to each experiment round 1 and round 2 oligo plates are annealed in a PCR machine by heating plates to 90°C and then decreasing the temperature by 1°C every 30 seconds until room temperature is reached.

#### Oligonucleotides for reverse transcription

To increase the barcoding space further, barcoded RT-primers are used. Eight pairs of RT-primers targeting the constant region of the TCR alpha and TCR beta locus were designed with a 4bp barcode and a 10bp UMI as well as a phosphorylated 5’ overhang complementary to the overhang of the round 1 top-strand barcoding oligo.

#### TCR-V-segment primer pool for multiplex PCR

Primers were initially designed by alignment of annotated C57BL/6J cDNA sequence (IMGT database) belonging to the same TCR-V-segment family. For each family 1-5 primers (depending on number and sequence similarity of TCR-V-segment families) with similar annealing temperature (+/- 1°C), length and G/C content were designed (see supplementary table X). Subsequently, C57BL/6J TCRα and TCRβ loci were aligned to the corresponding genomic sequence in the genomes of CAST/Ei, PWK/PhJ (evolutionarily closest publicly available genome compared to the used PWD/PhJ mouse strain) and SPRET/EiJ (genome data available as part of the Mouse Genome Project from Sanger Institute). Candidate primers were then BLAT searched against the aligned genomes to rule out the presence of SNPs in the primer binding region across all strains. All candidate primers were individually tested to exclusively amplify the corresponding V-segment(s) in reverse transcription reactions using RNA isolated from C57BL/6J CD8a^+^ T cells.

The final set of TCR-V-Segment primers consists of 58 individual primers (19 Vβ and 39Vα primers). Additional to the V-segment specific 3’ end of the primer, each primer also contains a common 5’ sequence used as target in the index-PCR. All V-segment primers were pooled at an equimolar ratio with a final concentration of 100 µM (1.72 µM of each primer). The primer pool was prepared once, and aliquots were frozen until used in an experiment to prevent biases introduced by varying primer pools across all experiments.

#### Cell fixation

After cell collection from tissue culture plates, 1ml of cell-suspension in PBS was added to 2.8ml of ice-cold PBS with 200µl of 16% PFA (ThermoFisher, 28908), for a final concentration of 0.8% PFA. After 10 minutes of incubation on ice 150µl 10% Triton-X was added to permeabilize cells and incubation on ice was continued for another 3 minutes. Cells were then centrifuged at 400xg 4°C for 5 minutes. Supernatant was discarded and the cell pellet was resuspended in 500µl 0.6M Tris-HCL pH8. Afterwards, 500µl of wash-buffer (PBS + 2% FBS and 0.4U/µl RNAseInhibitor (JenaBioscience, PCR-392L)) was added and cells were centrifuged at 400xg 4°C for 5 minutes. Washing was repeated once with 1ml wash-buffer before cells were counted and the concentration was adjusted to 50.000 cells/ml with wash-buffer.

#### Reverse transcription

10µl of fixed cells (∼50.000 cells) were added to each of 8 tubes of a prepared PCR-strip containing 1µl 10µM barcoded TCRalpha constant region RT-primer, 1µl 10µM barcoded TCRbeta constant region RT-primer, 7.5µl NEB TS Buffer (NEB, B0466SVIAL) and 2µl 10mM dNTPs (ThermoFisher, R0181). TCRalpha and TCRbeta RT-primers within each tube share the same tube-specific 4bp barcode. The number of reverse transcription reactions can be scaled up easily by increasing the number of prepared PCR strips. Typically, two PCRs strips for a total of 16 reverse transcription reactions were prepared resulting in a final cell count of ∼600.000 after barcoding (during the barcoding procedure about 25% of cells are lost due to repeated transferring and pooling of cells). Cells were then heated to 55°C for 5 minutes and rapidly cooled down to 4°C to allow pre-annealing of the RT-oligos to their target mRNAs. Afterwards, 6.3µl water, 1.5µl Maxima H Minus Reverse Transcriptase (ThermoFisher, EP0751) and 1ml RNAseInhibitor (JenaBioscience, PCR-392L) was added to each reaction for a final reaction volume of 30µl. Reverse transcription was carried out under the following conditions: 50°C for 10 minutes followed by 3 cycles of (8°C for 12 s, 15°C for 45 s, 20°C for 45 s, 30°C for 30 s 42°C for 2 minutes and 50°C for 3 minutes) and a final incubation at 50°C for 10 minutes. After reverse transcription cells were centrifuged at 400xg 4°C for 5 minutes. Supernatant was carefully discarded without disturbing the cell pellet. Cells were then resuspended in 50µl wash-buffer per tube and pooled in one 5ml tube and washing was repeated once.

#### Barcode ligation

All tubes used for pooling and washing of cells were coated with PBS +2% FBS to prevent cells from sticking to the plastic. Cells were resuspended in 2ml ligation buffer 1 (1460 µl water, 400 µl 10x T4 DNA ligase reaction buffer (NEB, B0202SVIAL), 100 µl T4 DNA ligase (NEB, M0202LVIAL), and 40 µl 10% Tween-20). 10µl of cell suspension was pipetted to each well of the two 96-well round 1 barcoding plates, taking care to not touch the liquid at the bottom of the plate. Plates were sealed with adhesive seals (ThermoFisher, AB0558) and incubated on a shaker for 40 minutes at room temperature. Afterwards, 3.5µl blocking oligo solution (20µM blocking oligo in water) was added to each well of both round 1 barcoding plates and incubation was continued for additional 20 minutes. The blocking oligo anneals to un-ligated round 1 top-strand oligos to prevent undesired ligations during the first cell pooling. Using a multichannel pipette, cells from both round 1 barcoding plates were pooled into a reservoir and then transferred to a 5 ml tube. Afterwards cells were centrifuged at 750xg 4°C for 3 minutes, supernatant was discarded, and cells were resuspended in 5ml of ligation buffer 2 (2260 µl water, 700 µl T4 DNA ligase reaction buffer, 100 µl T4 DNA ligase, 1900 µl annealing buffer and 40 µl 10% Tween-20). 25µl of cell suspension was pipetted into each well of the two 96-well round 2 barcoding plates, again without touching the liquid at the bottom of the wells. Plates were sealed and incubated for 40 minutes on a shaker at room temperature. Cells were then pooled as described before, centrifuged at 750xg 4°C for 3 minutes and resuspend in 200µl wash buffer. 1x DAPI (ThermoFisher, D1306) was added, and cells were counted on the Evos Countess II. The concentration of cells was adjusted to 2×10^6^ cells/ml and 5 µl of cell suspension was transferred to separate tubes of PCR-strips for the generation of sub-libraries. The number of cells in each sub-library determines the expected number of barcode collisions in each sub-library. The number of collisions can be calculated with the formular used in the birthday problem. Here the total number of barcodes B is 294.912 (8 reverse transcription barcodes * 192 round 1 * 192 round 2 ligation barcodes) with a cell count of N = 10.000 cells per sub-library. The number of expected barcode collisions therefore is:

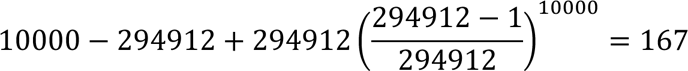

With 167 barcode collisions the expected collision rate is ∼1.67% in each sub-library.

#### Reverse Crosslinking

8 µl reverse crosslinking buffer (1% SDS, 100mM Tris-HCl pH8 and 100mM NaCl), 2 µl Proteinase K (Qiagen, RP107B-1) and 5 µl water was added to each tube with 5 µl sub-library for a final volume of 20 µl per reaction. Reverse crosslinking was done at 62°C for 2 hours on a shaker followed by a final incubation at 95°C for 15 minutes to inactivate Proteinase K. Afterwards, 12µl 10% Tween-20 was added to each sub-library to quench SDS before PCR.

#### cDNA library preparation

After reverse crosslinking and SDS quenching 48µl multiplex-PCR mix (23 µl water, 16 µl 5x Q5 reaction buffer, 3.2 µl TrueSeq-i7-long primer, 3 µl 10mM dNTPs, 2 µl 100 µM TCR-V-Segment primer pool and 0.8 µl Q5 DNA polymerase) was directly added to each sub-library for a final PCR reaction volume of 80µl. PCR was done using the following parameters: 98°C 2min, then 10 cycles of (98°C 20 s, 63°C 30 s, 72°C 2 minutes) and a final incubation at 72°C for 5 minutes. After PCR amplified cDNA was purified by bead clean-up using custom size-selection beads at a ratio of 1.2x beads to PCR reaction (100 µl beads) to get rid of excess primers from the multiplex PCR. During this clean-up it is important to not cross-contaminate different sub-libraries as they have not yet received their sub-library specific index.

14.5 µl index-PCR mix (10 µl 5x Q5 reaction buffer, 2 µl TrueSeq-i7-long primer, 2 µl 10mM dNTPs and 0.5 µl Q5 DNA polymerase) was added to each sub-library. Afterwards, 2.5µl of a unique 10µM Nextera N5xx primer was added to each sub-library for a final reaction volume of 50 µl. Index PCR was done using the following parameters: 98°C 2min, then 12 cycles of (98°C 20 s, 63°C 30 s, 72°C 2 minutes) and a final incubation at 72°C for 5 minutes. After index PCR sub-libraries were purified using 1.2x size-selection beads as described above. cDNA concentration of each sub-library was measured, and sub-libraries were then pooled at an equimolar ratio. Before freezing the pooled libraries until sequencing they were quantified using the Qubit HS dsDNA Quantification Kit and run on the Agilent 2100 bioanalyzer with a High Sensitivity DNA kit.

#### DNA size selection with custom beads

To prepare custom DNA size-selection beads, 750 µl of SPRIselect (Beckman Coulter, B23318) were transferred to a 1.5 ml tube and placed on a magnetic stand. Supernatant was discarded and beads were washed once with 1 ml Tris-HCl pH 8. Beads were then resuspended in 50 ml bead buffer (22 mM PEG-8000, 2.5 M NaCl, 10mM Tris HCl pH 8, 1 mM EDTA in water).

In general, size selection beads are added to the solution containing DNA at a defined ratio to bind DNA of a specific length (e.g., 1.2x beads will bind dsDNA >200bp). After binding DNA for 5 minutes, tubes are placed on a magnetic stand and supernatant is discarded (or transferred to a different tube in case of upper cut-off size selection). Beads are then washed twice with 80% EtOH before DNA is eluted from the beads by adding the desired volume of water or 10mM Tris HCl pH 8.

#### Sequencing

All TCR cDNA libraries have been sequenced on the Nova-seq 6000 platform by Illumina using S4 2×150bp v1.5 kits with the following sequencing-cycle set-up: Read1: 150 cycles, Index1: 17 cycles, Read2: 150 cycles and Index2: 8 cycles.

#### Cost of CITR-seq experiments

In CITR-seq all molecular reactions are carried out in bulk for ∼5.000-50.000 cells depending on the protocol step. This offers significant cost advantages, especially in contrast to plate-based single-cell protocols in which all molecular reactions are done separately for each cell. Enzymes needed for one experiment (using 500.000 input cells) in our hands cost about 350$ (ligase, reverse-transcriptase, polymerase, RNAse inhibitor etc.). The required barcoding oligos can be bought in high quantities and are then sufficient for many CITR-seq runs bringing down the oligo costs to less than 50$ per experiment. Collectively, the cost for library preparation in each experiment is therefore roughly 400$.

### Analysis

#### CITR-seq sequencing data pre-processing

Demultiplexing of fastq-files was done using a custom script, allowing one nucleotide mismatch in the cellular barcode sequence (relative to the barcode whitelist). Afterwards, adapter sequences were trimmed from the sequencing reads using *cutadapt* [67]. We then used *UMItools* [68] to extract the 4bp in-line barcode sequence from each sequencing read. For each read the in-line barcode and the barcode sequence extracted from the corresponding index reads were combined. The combined barcode sequences were then added to the 5’ end of read1. Afterwards, the full barcode information is present at the beginning of read 1 (16bp) followed by the UMI (10bp) and the 150bp sequencing read. Read 2 contains just the 150bp sequencing read. This pre-processing of sequencing reads modifies the fastq-files to be easily integrated into the subsequent MiXCR-pipeline.

#### Species-specific V(D)J reference libraries

To construct individual V(D)J reference libraries for PWD/PhJ, CAST and SPRET we built on the strategy used in the *findAlleles* function implemented in the MiXCR [47] software. First, we used full-length TCR sequencing data of each species generated using the 10x Genomics Immune Profiling Kit (see below), to assemble gene-segment candidate-alleles: Raw sequencing fastq files were processed using *Cellranger VDJ* supplying the built-in mm10 based VDJ-reference (GRCm38-ensemble-7.0.0). In this pipeline fragmented reads are combined into full length contigs based on sequence overlap in reads and matching cellular barcodes. We used the generated “filtered_contig.fastq” output and passed it directly to the MiXCR alignment step (“*align*”, --species mmu, -- preset generic-amplicon --floating-left-alignment-boundary --floating-right-alignment-boundary C --rna) to generate binary vdjca-files. We then used *mixcr exportAlignments* (--dont-impute-germline-on-export -allNFeatures UTR5Begin FR3End) to extract gene-features so that SNPs in candidate-alleles are not modified to match the provided reference. For each candidate V(D)J-allele we then used the extremely unique combination of associated UMI and CDR3 sequences to distinguish low-frequency alleles from alleles generated by sequencing or PCR errors by requiring each allele to be identified with at least two unique CDR3/UMI combinations. The list of identified V,D and J segment alleles was then used to generate a MiXCR compatible reference libraries for each species using the *buildLibrary* function implemented in MiXCR. Since the underlying RNA-based input libraries are generated using template-switching rather than multiplex-PCR, they allow for the discovery of *de novo* V(D)J-segments since template-switch based cDNA libraries do not require previous knowledge of the entire set of gene-segments for amplification.

#### Full list of V(D)J genes/families analyzed in cross-species comparisons

All names of V(D)J genes/families correspond to the official IMGT nomenclature [57]. Pseudogenes as well as extremely low-expressed genes (< 200 transcripts across all ∼5×10^6^ T cells of all species) are excluded from the analysis. *Trbv24* (all species) and *Trbv31* (PWD) were excluded from the analysis due to failure of amplification during the multiplex PCR. The remaining list contains the following V(D)J genes/families:

1. **Trav-families:** Trav1, Trav2, Trav3, Trav4, Trav5, Trav6, Trav7, Trav8, Trav9, Trav10, Trav11, Trav12, Trav13, Trav14, Trav15, Trav16, Trav17, Trav18, Trav19, Trav21
2. **Trbv-genes:** Trbv1, Trbv2, Trbv3, Trbv4, Trbv5, Trbv12-1, Trbv12-2, Trbv13-1, Trbv13-2, Trbv13-3, Trbv14, Trbv15, Trbv16, Trbv17, Trbv19, Trbv20, Trbv21, Trbv23, Trbv26, Trbv29, Trbv30, Trbv31
3. **Traj-genes:** Traj2, Traj4, Traj5, Traj6, Traj7, Traj9, Traj11, Traj12, Traj13, Traj15, Traj16, Traj17, Traj18, Traj21, Traj22, Traj23, Traj24, Traj26, Traj27, Traj28, Traj30, Traj31, Traj32, Traj33, Traj34, Traj35, Traj37, Traj38, Traj39, Traj40, Traj42, Traj43, Traj44, Traj45, Traj47, Traj48, Traj49, Traj50, Traj52, Traj53, Traj54, Traj56, Traj57, Traj58
4. **Trbj-genes:** Trbj1-1, Trbj1-2, Trbj1-3, Trbj1-4, Trbj1-5, Trbj1-6, Trbj2-1, Trbj2-2, Trbj2-3, Trbj2-4, Trbj2-5, Trbj2-16, Trbj2-7

All gene names in the generated species-specific V(D)J reference files correspond to the closest relative (by sequence identity) in mm10 based MiXCR reference library.

#### Alignment of sequencing reads using MiXCR

Sequencing reads in pre-processed fastq-format were integrated into a custom MiXCR pipeline (MiXCR version 4.5.0) using the following steps:

1. *mixcr align* `-- preset generic-ht-single-cell-amplicon-with-umi` `-- library` Species Specific custom library (see above) `-- tag-pattern ^(CELL:N(16))(UMI:N(10))(R1:*)\^(R2:*)` `-- floating-left-alignment-boundary` `-- floating-right-alignment-boundary C` `- OvParameters.geneFeatureToAlign=VRegionWithP` `- OminSumScore=100`
2. *mixcr refineTagsAndSort*
3. *mixcr assemble* `-- assemble-clonotypes-by CDR3` `-- cell-level`

We then used *mixcr exportClones* to extract the required information for all downstream analysis (e.g., cellular barcodes, transcript counts, V(D)J segments, CDR3 amino acid and nucleotide sequence etc.).

#### Construction of 10x Genomics Single Cell Immune Profiling sequencing libraries

We generated four sequencing libraries (from 10-week-old male mice, primary CD8^+^ T cells of one of each: BL6, PWD, CAST, SPRET, see cell isolation described above) using the 10x Genomics Immune Profiling platform (Chromium Next GEM Single Cell 5’ Kit v2) according to the manufacturer’s instructions. T cells from each mouse were used in two separate reactions, each with 2.500 input cells (eight total reactions). V(D)J sequencing libraries were sequenced at 5.000 reads/cell. Raw sequencing data was pre-processed as described above and then aligned to species-specific V(D)J references using the outlined MiXCR pipeline.

#### Assignment of parental alleles in F1 hybrids

Pre-processed fastq-files of all F1 hybrid samples were aligned using MiXCR as described above. Importantly, the F1 hybrid samples were aligned to both parental V(D)J references and the alignment scores for V- and J-genes were extracted (*mixcr exportAlignments -vHitScore* and *-jHitScore*). We then compared the alignment scores for V- and J-genes from both alignments for each sequencing read. Each gene segment was then assigned to one parental species based on the higher alignment score in both alignments. Absence of SNPs in a gene-segment lead to identical alignment scores and therefore the respective reads were only assigned to a parental allele if the second gene segment in the same read was assigned to one parental allele. Reads in which both V- and J-segments had identical alignment scores in both alignments (e.g. no parental SNPs in both gene segments) as well as reads in which V- and J-parental assignment disagreed were discarded from the analysis together with all other reads sharing the respective identical cellular barcode.

#### Comparison of CITR-seq data with publicly available datasets from Parse Bioscience and 10x Genomics

Absolute counts of paired αβ-TCRs shown in **Fig. 2D** were taken from the following datasets:

1. **Parse Bioscience [44]:** TCR Sequencing of 1 Million Primary Human T Cells in a Single Experiment (primary human Pan T cells, sequencing depth: 5000 reads/cell).
2. **10x Genomics Single Cell Immune Profiling [45]:** CD8+ T cells of Healthy Donor 2 (v1, 150×91), Single Cell Immune Profiling Dataset by Cell Ranger v3.0.2, 10x Genomics, (2019, May, 9)

The UMI/cell recovery rates in CITR-seq were compared to the UMI/cell recovery rates in two publicly available datasets provided by Parse Bioscience:

1. **Parse Bioscience [44]:** TCR Sequencing of 1 Million Primary Human T Cells in a Single Experiment (primary human Pan T cells, sequencing depth: 5000 reads/cell)
2. **Parse Bioscience [46]:** Performance of Evercode TCR in Activated Human T cells (Pan T cells after 72h activation using CD3/CD28 beads + IL-2 supplementation, sequencing depth: 5000 reads/cell)

Datasets are available from their website (https://www.parsebiosciences.com) and the specific UMI/cell rates were extracted from the “TCR:Barcode Report (TSV)” tables (column: “transcript_count”)

#### PCA of VJ-pairing and central CDR3 4mer abundance

We conducted Principal Component Analysis (PCA) on two different datasets.

##### 1) V-J pairing

The first PCA was done to compare total counts of observed V-J pairs across all sample down-sampled to a common cell count of 5.000 T cells (each with one associated TCRα and TCRβ chain). The count-tables were analyzed using DESeq2’s[69] *varianceStabilizingTransformation* (*vst*, blind=FALSE, nsub=300) and PCA was conducted on the top 300 most variable V-J pairs (*plotPCA*, ntop=300). Using this parameters PC1-4 explain approximately 67% of variance in V-J usage across samples.

##### 2) CDR3 4mers

The second PCA analysis was done on a set of amino acid 4mers (or 4mer pairs) extracted from the central region of CDR3 amino acid motifs (the three most 3’ and 5’ amino acids were trimmed from the motif). Initially three randomly chosen 4mers were extracted from each trimmed CDR3 motif. For paired CDR3αβ motifs we extracted 3 random 4mer pairs of the respective CDR3α and CDR3β motifs associated with a cellular barcode. Subsequently, we generated count-matrices with the total counts of each 4mer across the 32 individuals. We filtered the matrices to only contain 4mer/pairs that were observed at least once across three individuals of a specific genotype (final 4mer counts: 39.843 CDR3α, 56.538 CDR3β and 1.201.646 CDR3αβ 4mers). The filtered count matrices were then analyzed following the standard DESeq2 [69] workflow for un-normalized count-matrix inputs. Afterwards, PCA was done on the top 5000 most variable 4mers using the *plotPCA* function.

#### Diversity and overlap indices used for repertoire comparison in CITR-seq data

Many of the commonly used indices from the analysis of TCR repertoires within and across different samples, were originally developed to quantify the diversity of species in an ecosystem. For this reason, they are often classified as either *alpha-diversity* indices that measure species richness and/or evenness within a particular population or alternatively as *beta-diversity* indices, which evaluate differences or overlaps between different populations. Similar to species diversity studies in ecology, TCR repertoire diversity estimates suffer from inherent incompleteness of the sampled diversity, a problem first described as the unseen species problem [70]. The use of diversity indices for adaptive immune receptor analysis is reviewed here [71]. The following indices were used in this study:

1. **Shannon diversity index** [72] (for proportions) The Shannon diversity index considers both, species richness and species evenness to evaluate the entropy within a distribution of species:

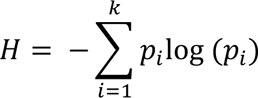

With *p_i_* = the proportion (frequency) in the group *k* (e.g. gene segments). The index can be normalized by dividing it by the maximum diversity. Which then is the normalized Shannon diversity index (nSDI) used in this study:

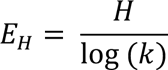

In the context of gene segment usage in nSDI of 1 would indicate that all gene segments are used at identical frequencies in a TCR repertoire.
2. **Jaccard index** The Jaccard index was developed by Paul Jaccard in 1901 and is commonly used to calculate the overlap (of CDR3 motifs) between two samples of TCRs:

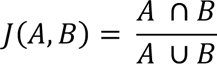

The Jaccard index calculates the intersection size divided by union size of two samples (A and B).

#### Classification of relative V(D)J gene usage in parental lines and their F1 hybrids

In classical F1 hybrid experiments, genes are often categorized into additive, dominant, over- and under-dominant, based on their expression in F1 hybrids relative to the parental individuals [73]. When adopted to V(D)J-gene usage in F1 hybrids and their parental lines it is important to note, that the frequency of a particular gene does not only depend on differences in gene expression regulation but is also influenced by biases during V(D)J recombination and thymic selection. We see that thymic selection introduces significant changes to V(D)J gene usage and therefore amplifies gene usage differences across species relative to the differences emerging from differential gene regulation alone. This effect is especially strong in F1 hybrids were almost all V(D)J genes show significantly different frequencies relative to the parental species (**Fig S5A**). Instead of using a *p*-value based classification, we therefore decided to rather compare the relative frequencies of V(D)J gene usage across F1 hybrids and the parental species. Accordingly, V(D)J gene frequencies are classified using the following criteria:

1. **Conserved:** Gene frequency in the F1 hybrid is within 1% of the frequency in both parents
2. **Dominant:** Gene frequency in the F1 hybrid is within 1% of the frequency in one parent and more than 1% larger or smaller than the frequency in the other parent.
3. **Additive:** Gene frequency in the F1 hybrid is more than 1% smaller than the frequency in one parent and more than 1% larger than the frequency in the other parent.
4. **Over-dominant:** Gene frequency in the F1 hybrid is more than 1% larger than the frequency in bother parents.
5. **Under-dominant:** Gene frequency in the F1 hybrid is more than 1% smaller than the frequency in bother parents.

## Acknowledgements

We thank all past and present members of the Chan and Jones laboratory for input into experimental design, helpful discussion and improving the manuscript. We especially thank Felicity Jones for her scientific input throughout the entire study. We thank Sinja Mattes and the remaining team of animal caretakers at the Friedrich-Miescher Laboratory led by Cemal Yilmaz. We thank Aurora Panzera and Christian Feldhaus from the Bio Optics core facility of the Max Planck Institute for Biology for their support with all FACS experiments. We thank Insa Hirschberg for supporting tissue culture experiments. We also thank the Genome Center of the Max Planck Institute for Biology for providing support with CITR-seq library sequencing. M.P. and D.S. are supported by an International Max Planck Research School fellowship. M.K. and Y.F.C are supported by the European Research Council Starting Grant 639096 “HybridMix” and Proof-of-Concept Grant 101069216 “Haplotagging”. The research done in this study is supported by the Max Planck Society.

## Author Contributions

M.P. and Y.F.C. designed the experiments. M.P. and V.S. developed the barcoding framework for CITR-seq. M.P. developed the rest of the protocol and performed all experiments. M.P. performed the computational analysis advised by Y.F.C. M.P. wrote the manuscript. V.S., D.S., M.K. and Y.F.C. provided support for the experiments and the computational analysis. All authors reviewed the manuscript. Y.F.C. direct the study.

## Supplement

**Supplementary Figure 1:**
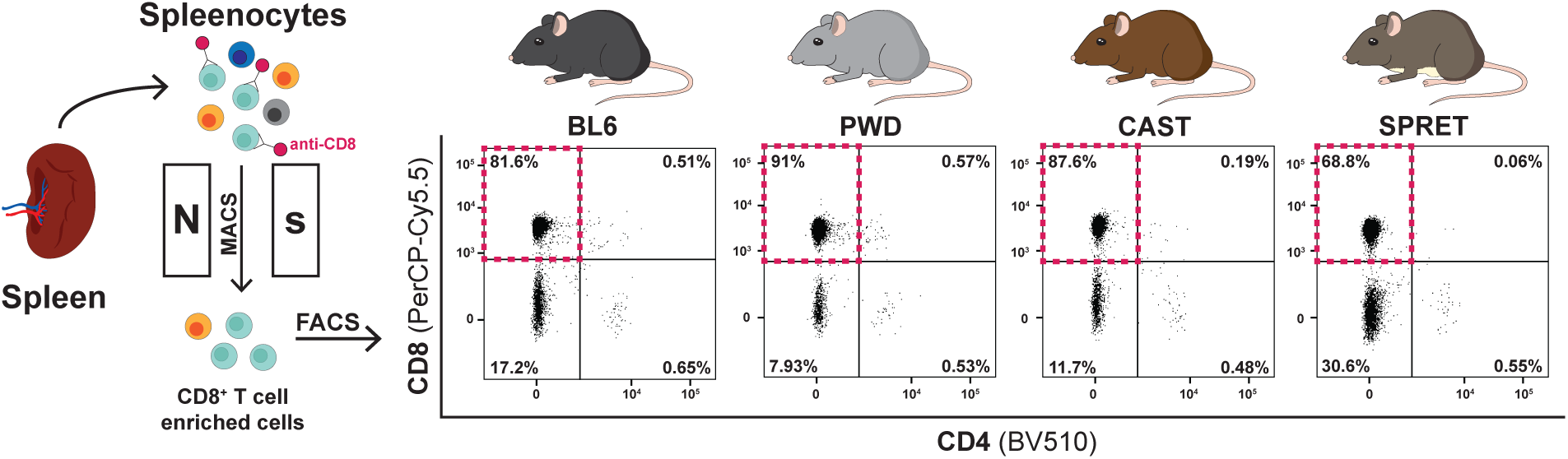
CD8^+^ T cells isolation strategy for all CITR-seq samples. **(A)** Spleenocyte cell-suspensions were pre-enriched for CD8^+^ T cells by magnetic extraction of anti-CD8 labeled cells (magnetic-activated cell sorting, MACS using Dynabeads™ FlowComp™ Mouse CD8 Kit). Afterwards pre-enriched cell-suspension was further purified using fluorescence activated cell sorting (FACS). Percentages in each quadrant of the FACS plots represent the mean frequencies of the respective cell population in the pre-enriched cell-suspension. CD8^+^ T cells in the top left quadrant (red box) were sorted and used for CITR-seq experiments.

**Supplementary Figure 2:**
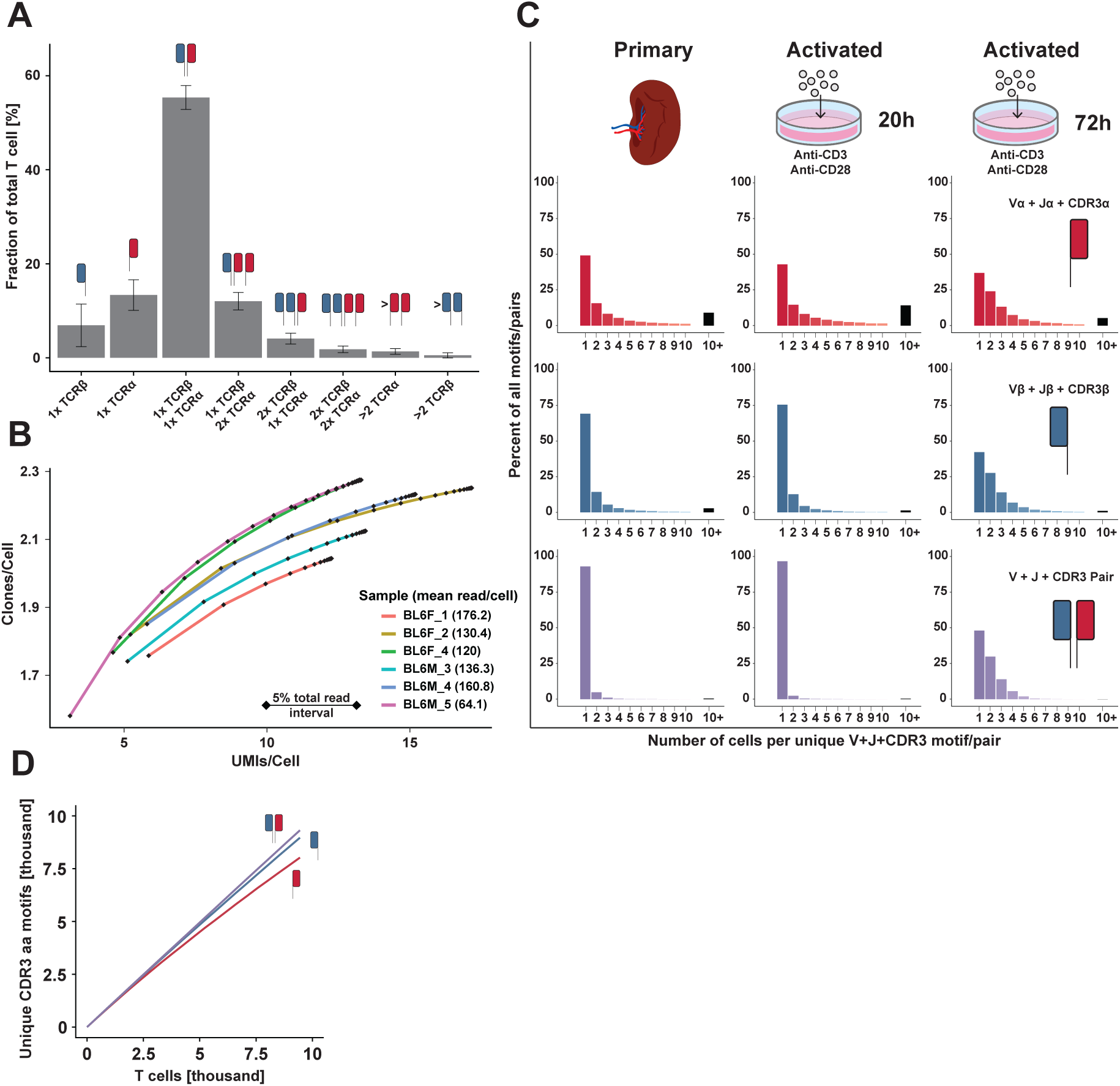
Additional analysis for CITR-seq validation. **(A)** Mean fraction of T cells assigned to different numbers of distinct TCRα and TCRβ chains (error bars represent the standard deviation across all 32 CITR-seq samples). Most T cells (∼55%) are associated with a single TCRα and a single TCRβ chain. Few T cells are associated with more than two TCRα (∼1.4%) or TCRβ (∼0.5%) chains, likely representing cell doublets or barcode collisions. **(B)** Saturation curve showing UMI/cell and clone/cell counts relative to the fraction of total sequencing reads. Diamonds represent the respective UMI/cell and clone/cell counts at intervals of 5% of sequencing reads (5% - 100% of reads) for six representative CITR-seq samples (all BL6 samples). The mean reads per cell are shown for the representative samples. **(C)** Clone size distributions (number of cells observed with a unique V+J+CDR3 TCR) in samples from primary T cells (left), 20h activated T cells (middle) and 72h activated T cells (right). The respective clone size distributions are shown for Vα+Jα+CDR3α TCRs (top), Vβ+Jβ+CDR3β TCRs (middle) or V+J+CDR3 paired αβ-TCRs (bottom). In contrast to primary and 20h activated T cells, 72h activated T cells show an increased clone size distribution caused by the onset of clonal expansion by prolonged T cell activation. **(D)** Total number of unique CDR3α, CDR3β or paired CDR3αβ amino acid motifs relative to the number of T cells across all 4 samples generated using the 10x Genomics Single Cell Immune Profiling Kit.

**Supplementary Figure 3:**
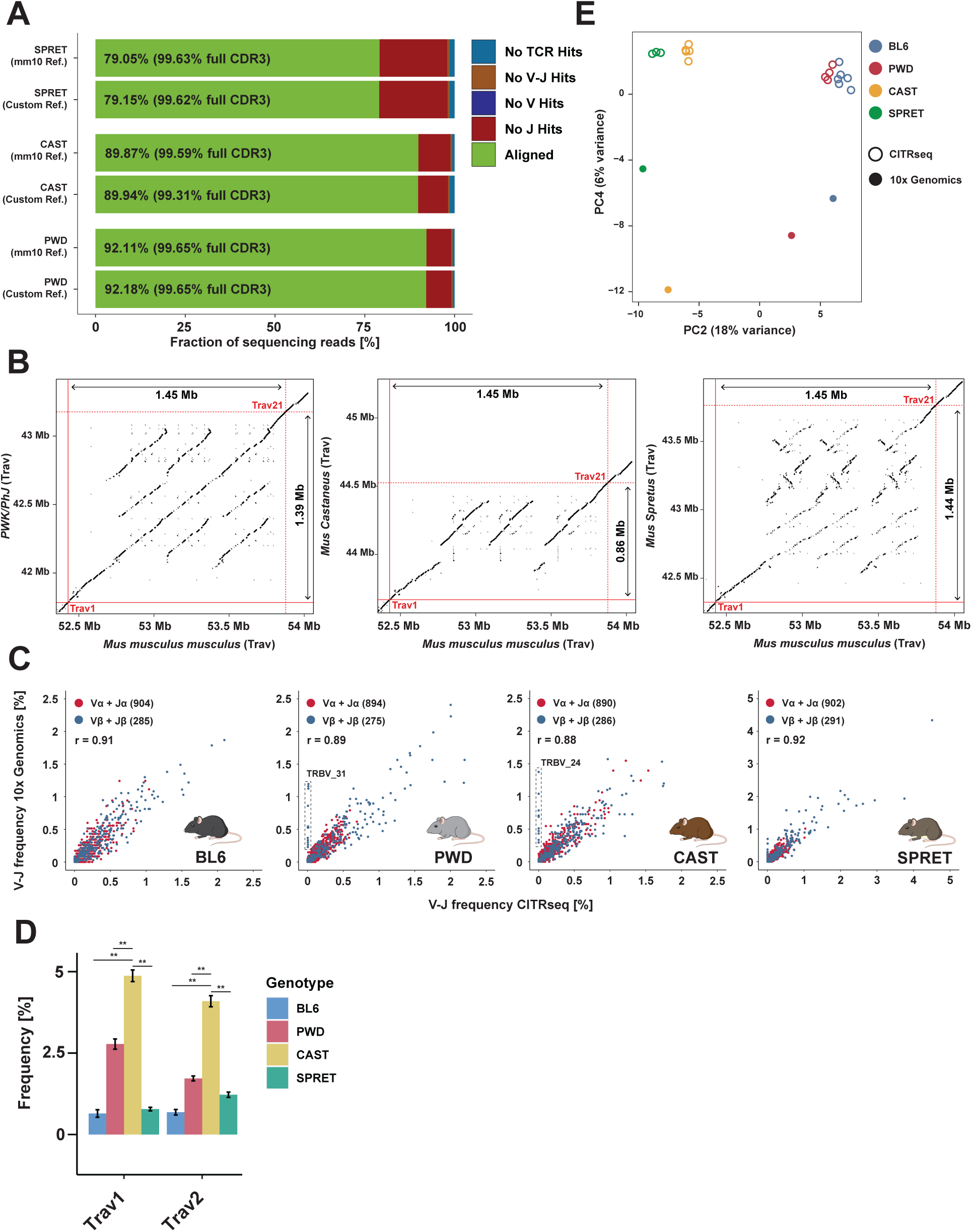
Different structure of TCRα loci across inbred species and comparison of observed V-J usage frequencies in different methods. **(A)** Fraction of sequencing reads that were successfully aligned to V(D)J genes using different reference libraries (green bars; full CDR3 coverage in brackets). Each stacked bar shows the mapping percentage for a representative sample from SPRET, CAST and PWD when aligned to the in-build mm10 based MiXCR V(D)J reference (top) and the species-specific custom V(D)J reference (see methods). All other colors in the stacked bar represent the reason for the failure of alignment. In all cases, the total fraction of successfully aligned reads is higher when using the species-specific custom library. **(B)** Dot plots of local alignment of genomic sequence from the GRCm38/mm10 TCR Vα locus to the PWK/PhJ (closest available genomic sequence to PWD, left), CAST (middle) and SPRET (right) genomic sequence of the TCR Vα locus. Intersections of the red lines indicate the location of the most distal (*Trav1*) and proximal (*Trav21*, dashed line) Vα genes. The genomic distances between these two Vα genes are shown. The central region of the Vα cluster is triplicated in BL6, PWK and SPRET relative to CAST. **(C)** Dot plots showing the mean frequency of single-chain V-J pairing in TCRα (red) and TCRβ (blue) chains observed in samples generated with -seq and 10x Genomics Single Cell Immune Profiling. The respective frequencies are shown for BL6, PWD, CAST and SPRET samples. Pearson-correlation and the total number of detected Vα-Jα and Vβ-Jβ are shown. Boxes highlight Vβ genes that are almost exclusively observed in 10x Genomics samples indicating failure of amplification for these Vβ genes by the multiple PCR primer pool used in CITR-seq. The respective Vβ genes were excluded from the analysis. **(D)** Usage frequencies of distal (5’) Vα genes (*Trav1*, *Trav2)* with one-to-one orthology across all four species. CAST mice have significantly higher frequencies of both genes compared to all other species (chi-squared test, ** *P-value* < 0.01). **(E)** PCA of combined Vα-Jα and Vβ-Jβ gene segment usage frequencies across all species as observed in samples generated with CITR-seq (empty circle) or 10x Genomics Single Cell Immune Profiling (filled circles). PC2 and PC4 are shown. PC4 contains 6% of the total variance across samples and separates samples by the respective methods used to generate the data.

**Supplementary Figure 4:**
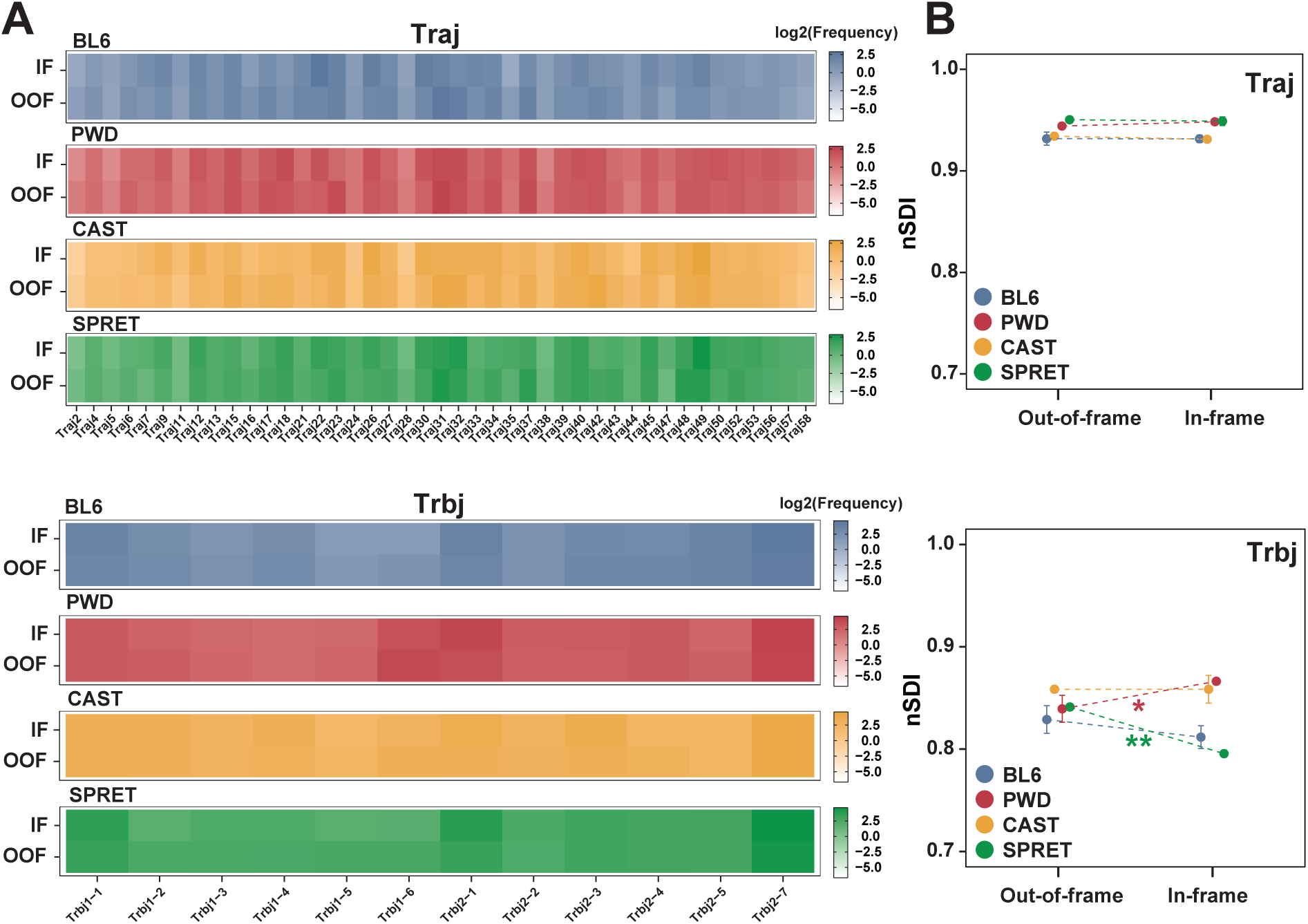
Comparison of Jα and Jβ gene usage in in-frame and out-of-frame TCRs. **(A)** Jα family (top) and Jβ gene (bottom) usage frequency (log_2_) heatmaps. Heatmaps show the mean intra-species J-usage in in-frame (IF) and out-of-frame (OOF) TCRs across all T cells. **(B)** Mean intra-species entropy in Jα-usage (top) and Jβ-usage (bottom) distributions calculated using the normalized Shannon diversity index (nSDI) for OOF and IF TCRs (error bars indicate the standard deviation in species replicates, significance calculated using paired t-test, * *P-value* < 0.05, ** *P-value* < 0.01).

**Supplementary Figure 5:**
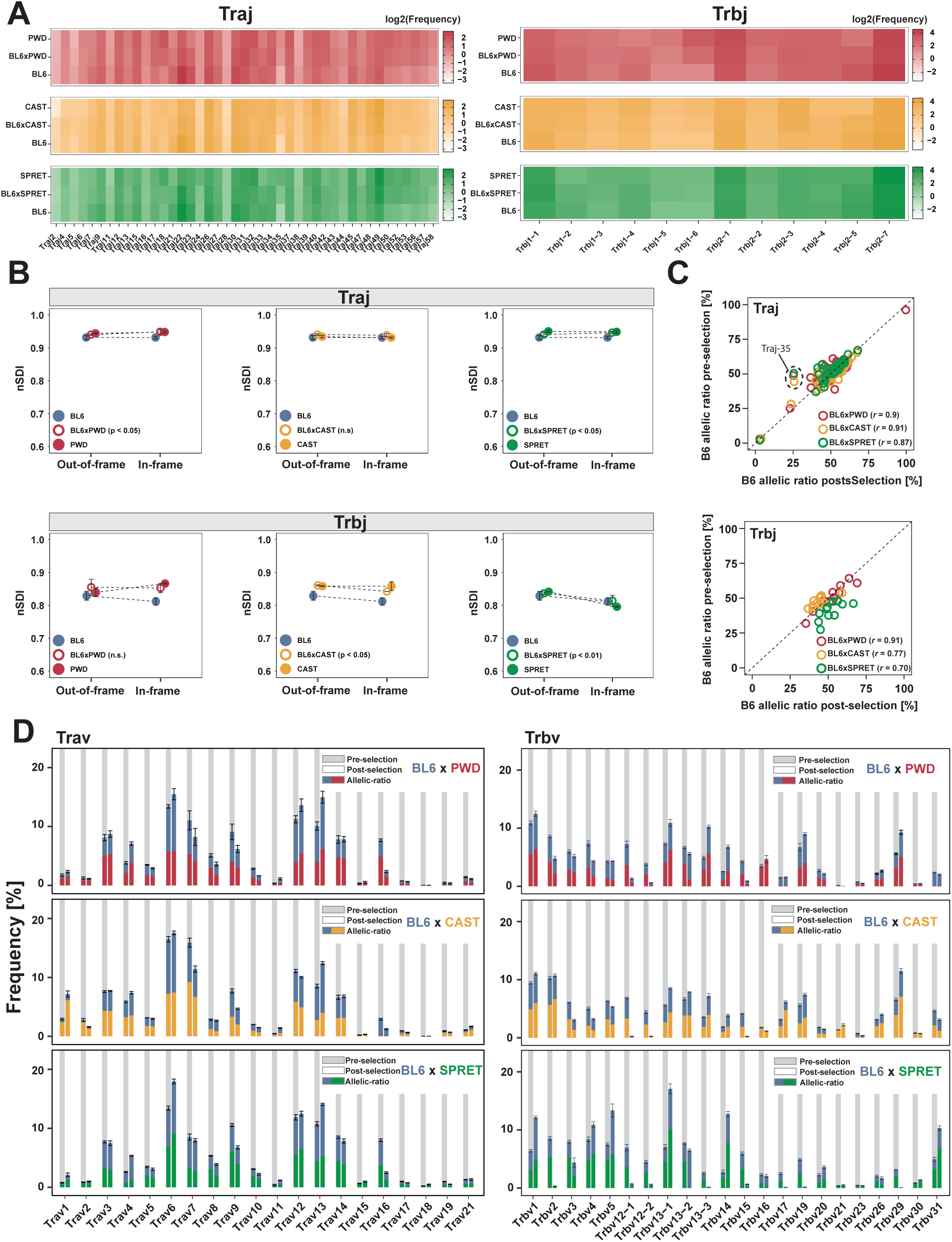
Usage frequencies of Jα and Jβ genes in F1 hybrids and the impact of thymic selection on their abundance. **(A)** Jα gene (left) and Jβ gene (right) usage frequency (log_2_) heatmaps of in-frame TCRs in F1 hybrids and their respective parental species. **(B)** Comparison of entropy of J-usage distribution in F1 hybrids and the respective parental species calculated using the normalized Shannon diversity index (nSDI) for OOF (left) and IF (right) TCRs (error bars indicate the standard deviation in species replicates, significance tested for F1 hybrid IF vs OOF contrast using paired t-tests). **(C)** Analysis of biased J gene allele usage in F1 hybrids. Plots show the percentage of BL6 Jα gene alleles and Jβ gene alleles in post- (x-axis) and pre-selection (y-axis) TCRs. Each circle represents a Jα-gene (top) or Jβ-gene (bottom). Pearson-correlation was calculated for post- and pre-selection J gene usage. Genes with substantial changes in allelic ratios in pre- and post-selection repertoires are highlighted (*Traj35*). **(D)** Detailed representation of the mean Vα-family and Vβ-gene usage frequencies in F1 hybrids in pre-selection (grey background) and post-selection (white background) TCRs. Stacked bars show the allelic ratio in the respective V-gene/family (error bars indicate the standard deviation in species replicates).

**Supplementary Figure 6:**
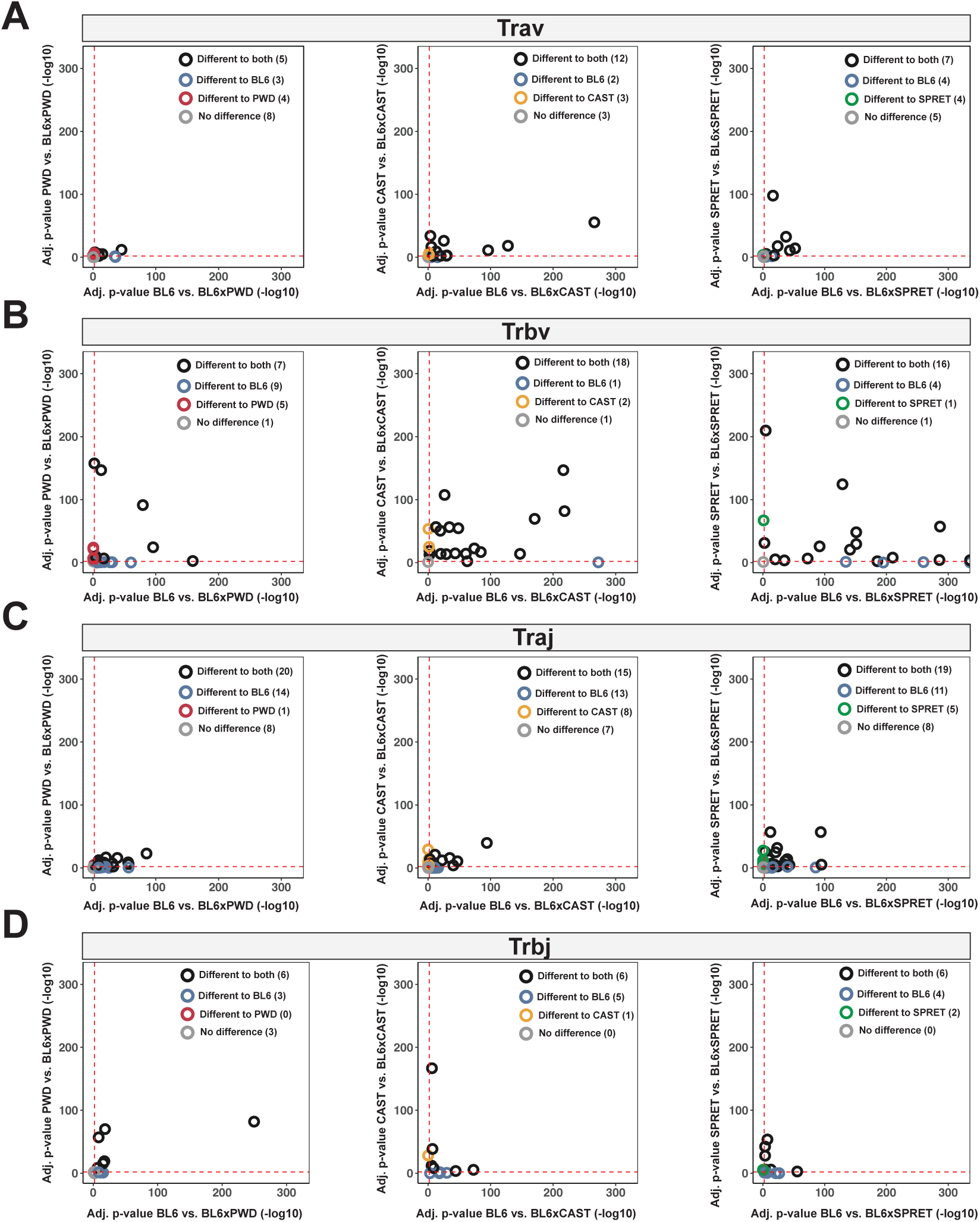
*P-values* (-log_10_) of V/J gene usage frequency changes in F1 hybrids relative to both parents. Adjusted *P-values* for V and J gene frequency changes in F1 hybrids relative to their parents. Plots show *P-values* for BL6xPWD (left), BL6xCAST (middle) and BL6xSPRET (right) for Vα-families **(A)**, Vβ-genes **(B)**, Jα-genes **(C)** and Jβ-genes **(D)**. *P-values* have been calculated for differences in absolute count of TCRs with the respective V/J genes using Wald-test. Dashed red lines show that *P-value* cut-off of *P* < 0.01. Genes with significant (*P* < 0.01) changes relative to both parents (empty black circles), to the BL6 parent (empty blue circles), the respective other parent (PWD, CAST, SPRET, empty red, yellow, green circles) as well as genes with no differences to both parents (empty grey circles) are shown with the respective total counts of genes in each category.

**Supplementary Figure 7:**
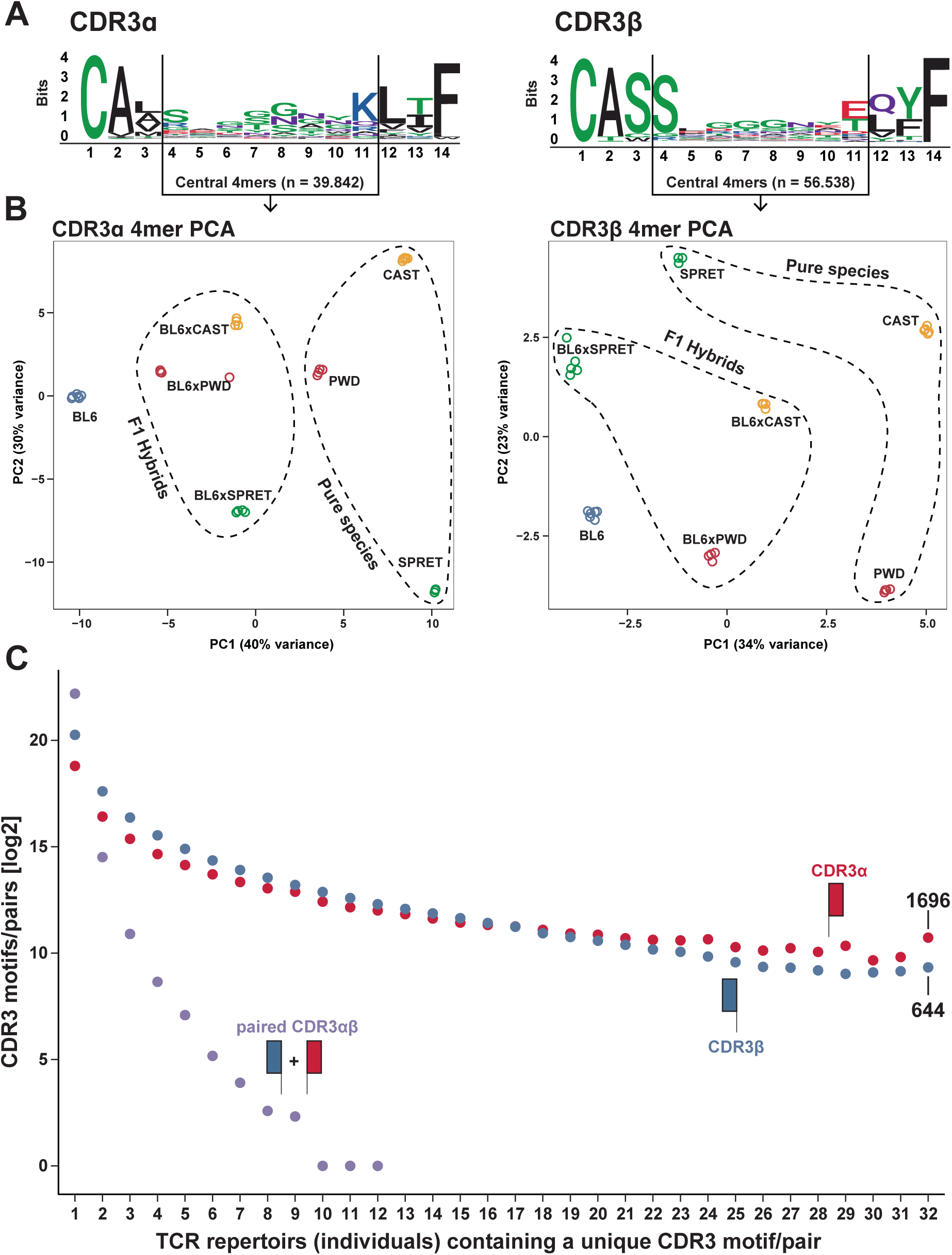
Comparison of single-chain CDR3α and CDR3β amino acid motifs in all 32 CITR-seq samples. **(A)** Positional diversity of amino acids in 10,000 randomly chosen CDR3α (left) and CDR3β (right) single-chain motifs from all 32 CITRs-eq samples. Black lines mark the central region of CDR3 motifs. Three 4mers were randomly selected from this central region of all CDR3 sequences across the 32 individuals. The filtered count-matrix of 4mers (see methods) contained 39,842 unique CDR3α and 56,538 unique CDR3β 4mers. **(B)** PCA analysis done using the 4mer count-matrices of CDR3α 4mers (left) and CDR3β 4mers (right). PC1 and PC2 are shown. 4mer samples cluster by genotype of the underlying sample. **(C)** Number of TCR repertoires (of individuals) in which each unique CDR3α (red), CDR3β (blue) or paired CDR3αβ (purple) motif is observed. 1,696 CDR3α motifs and 644 CDR3β motifs have been observed in TCR repertoires of every single individual analyzed in this study. Identical CDR3αβ motifs have not been observed in more than 12 individuals.

**Supplementary Table 1:**
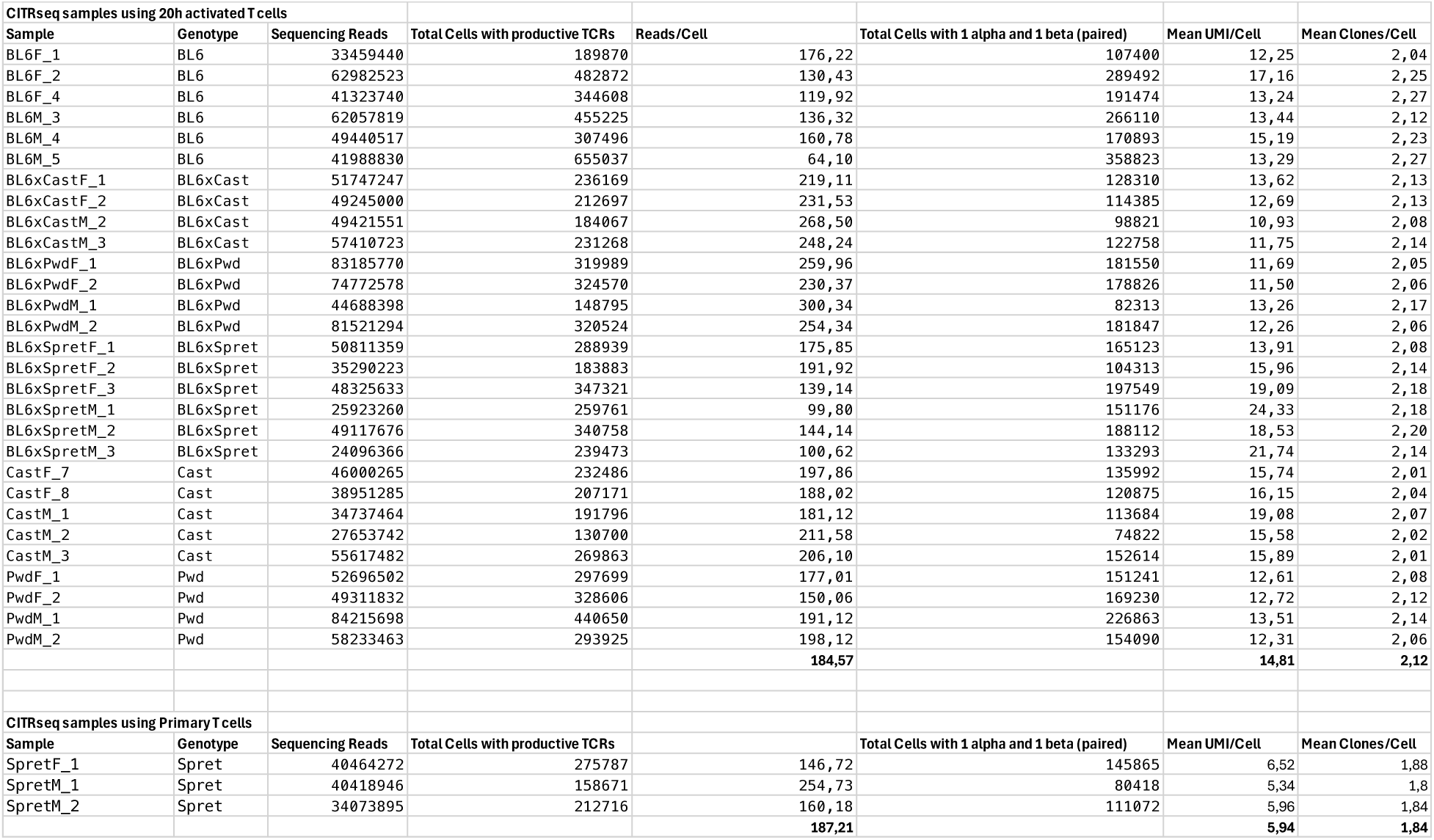
Detailed sample list of all 32 CITR-seq samples analyzed in the present study.

**Supplementary Table 2:**
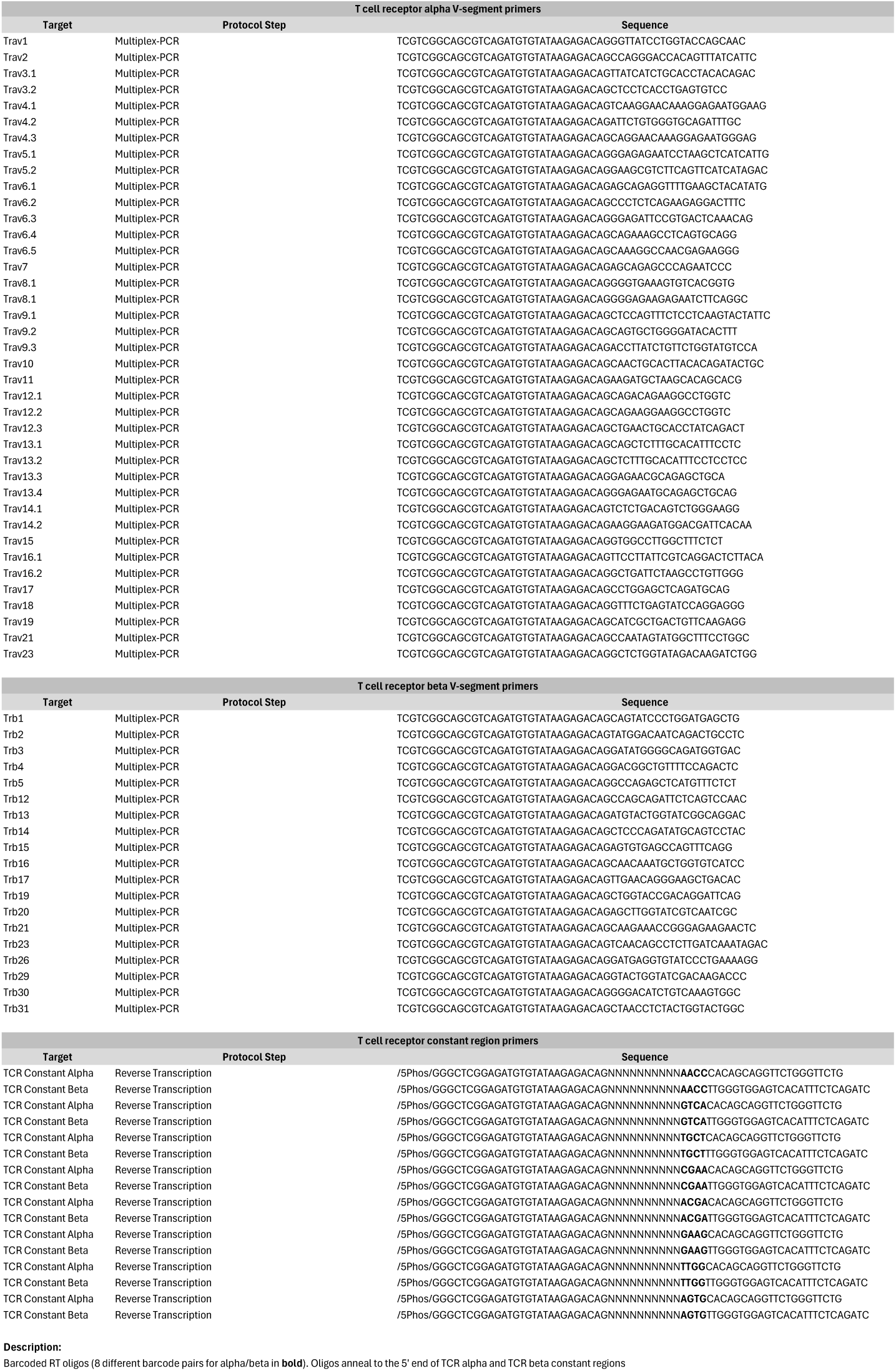
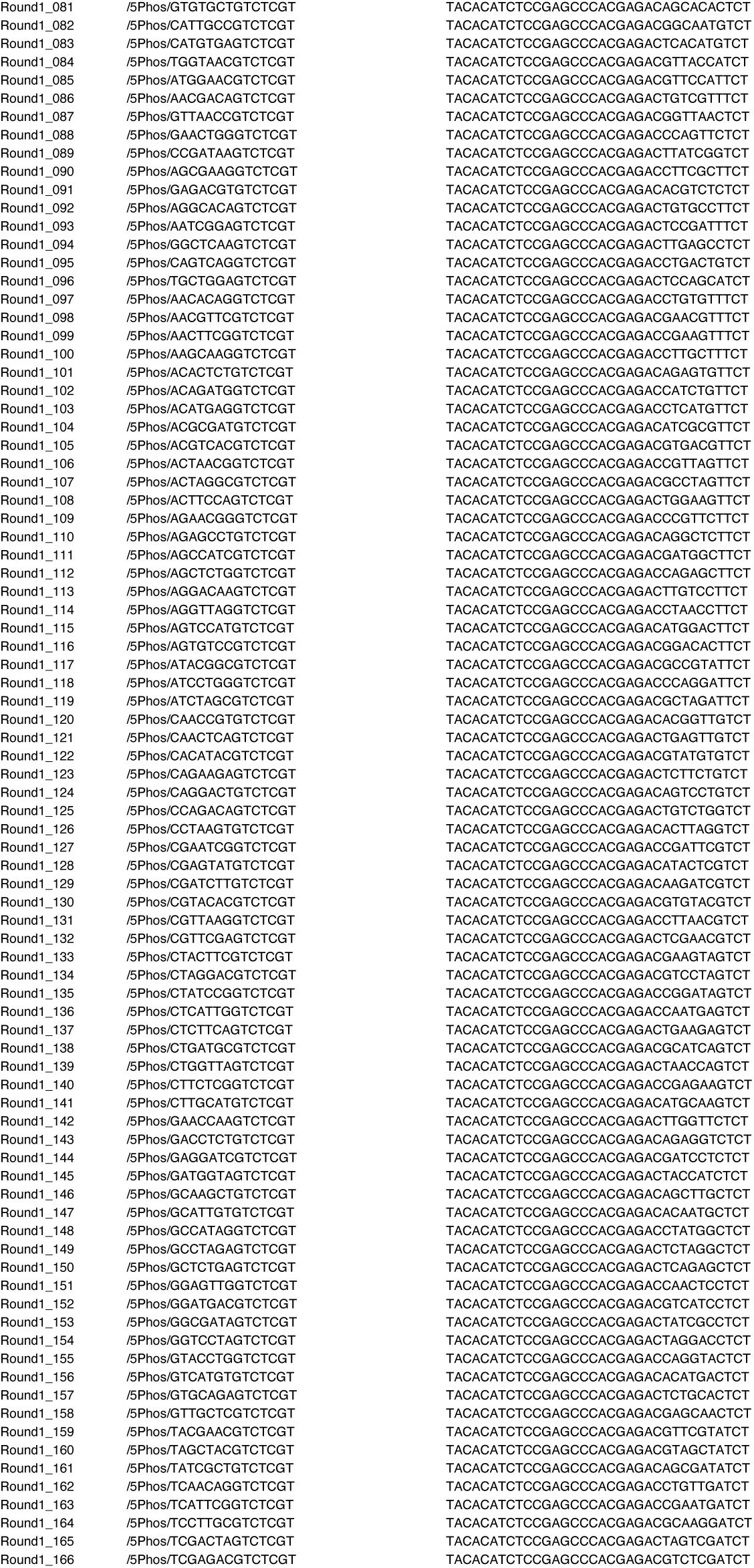
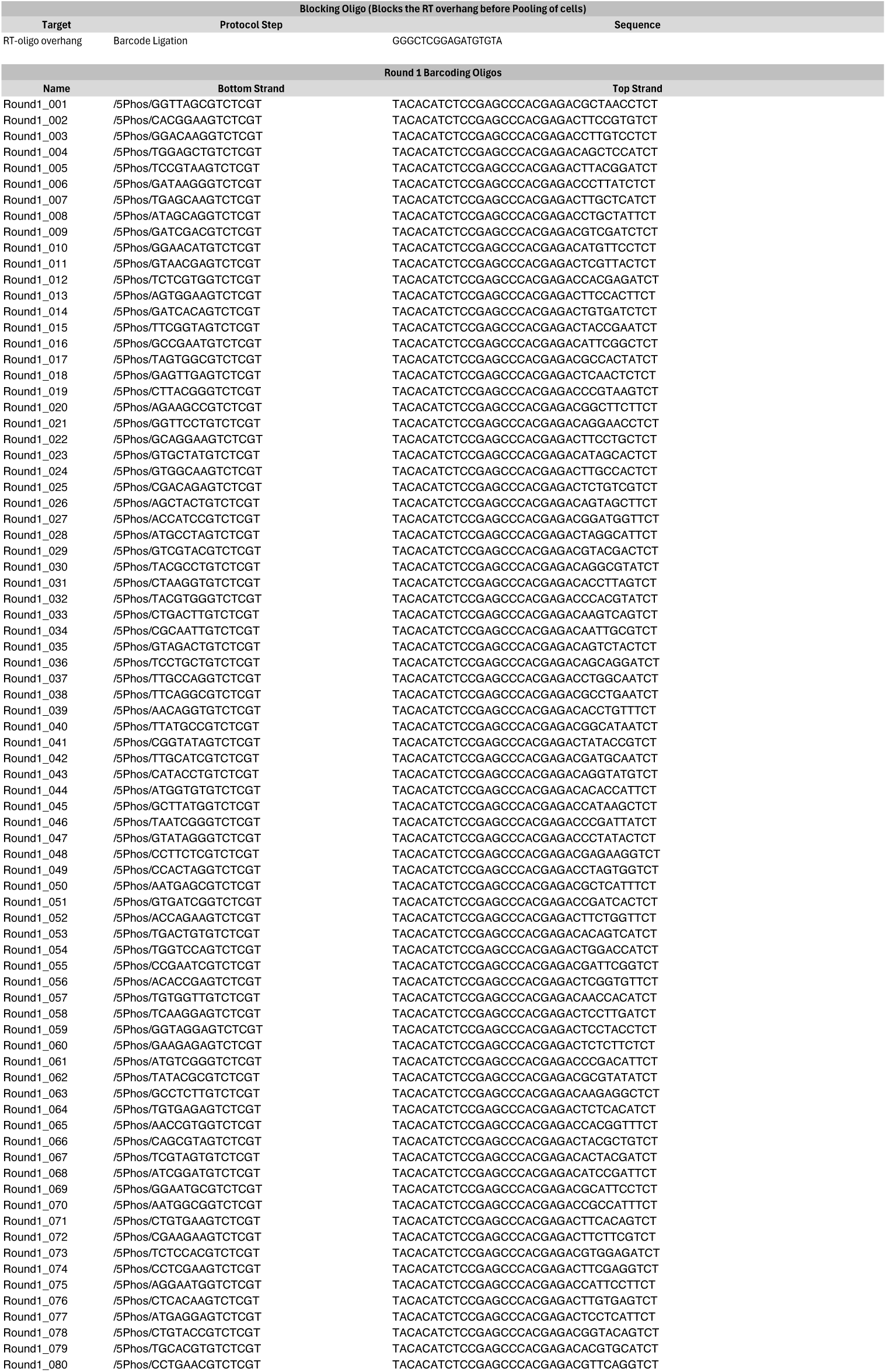
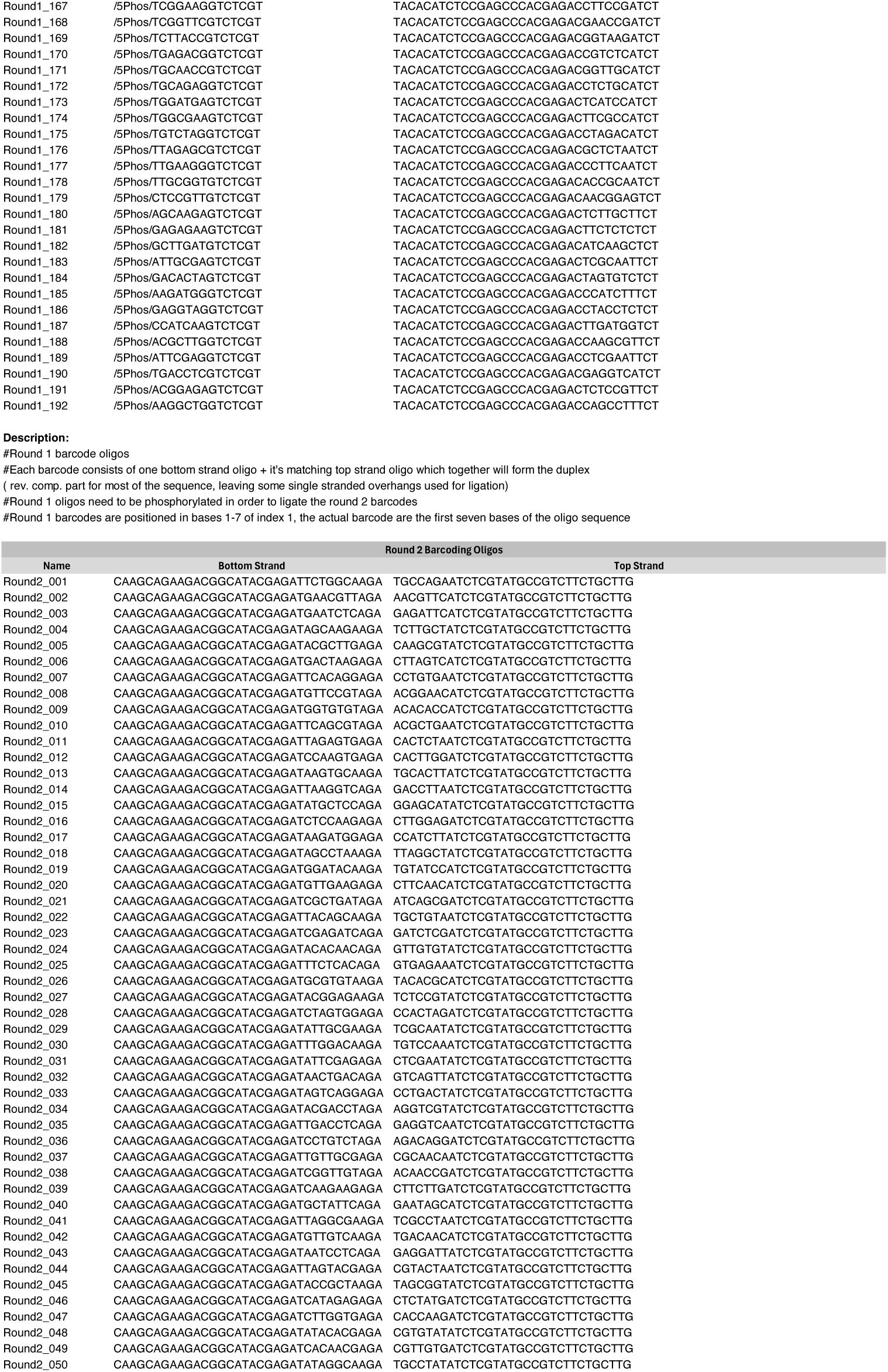
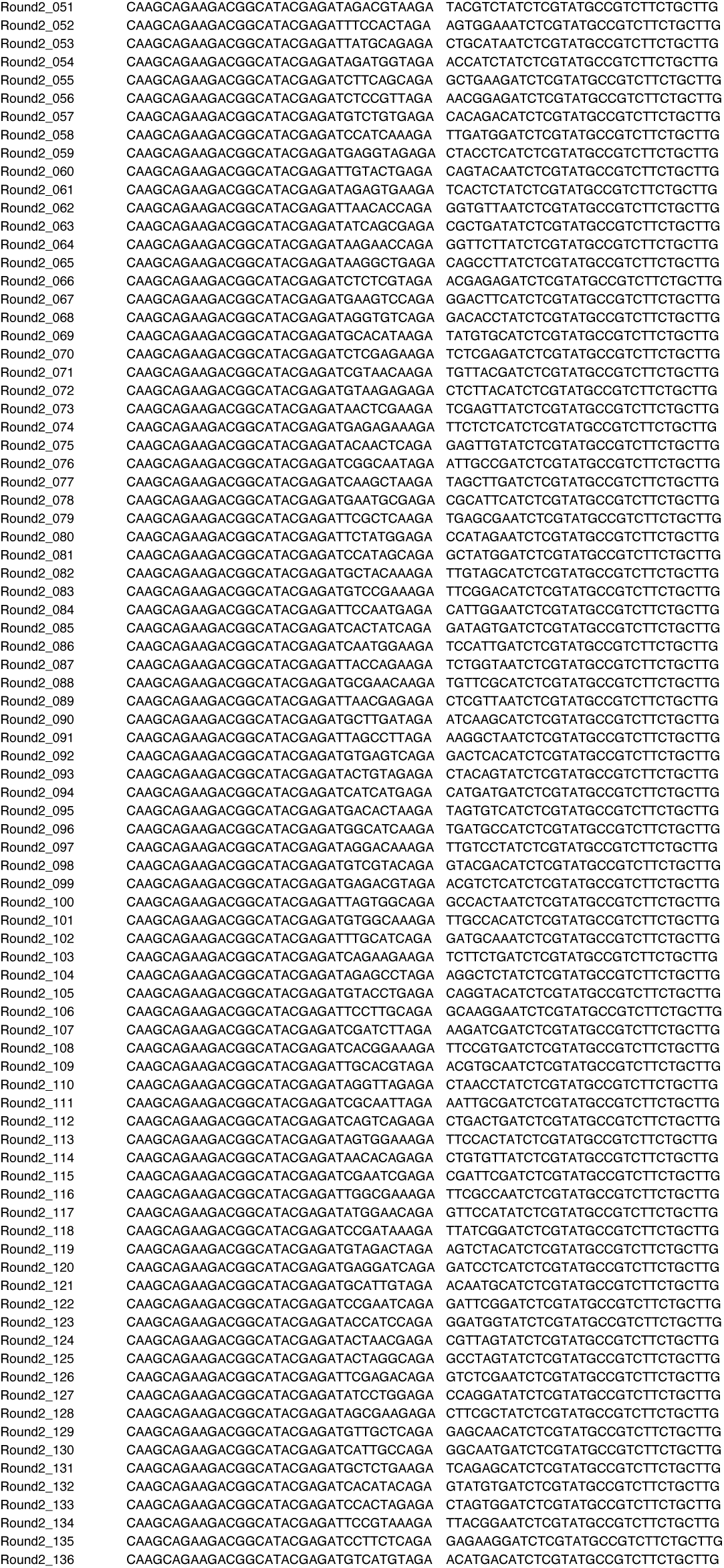
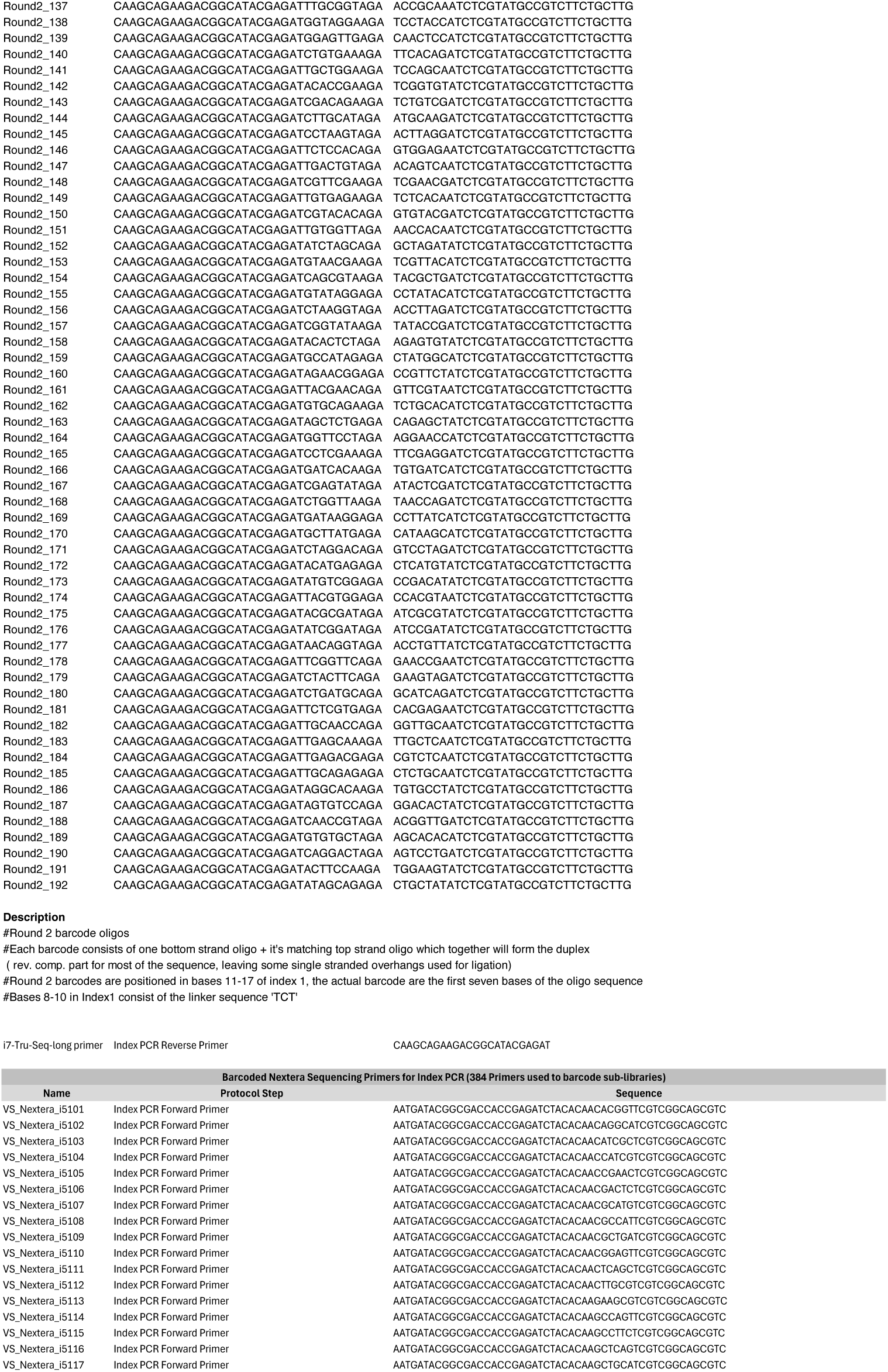
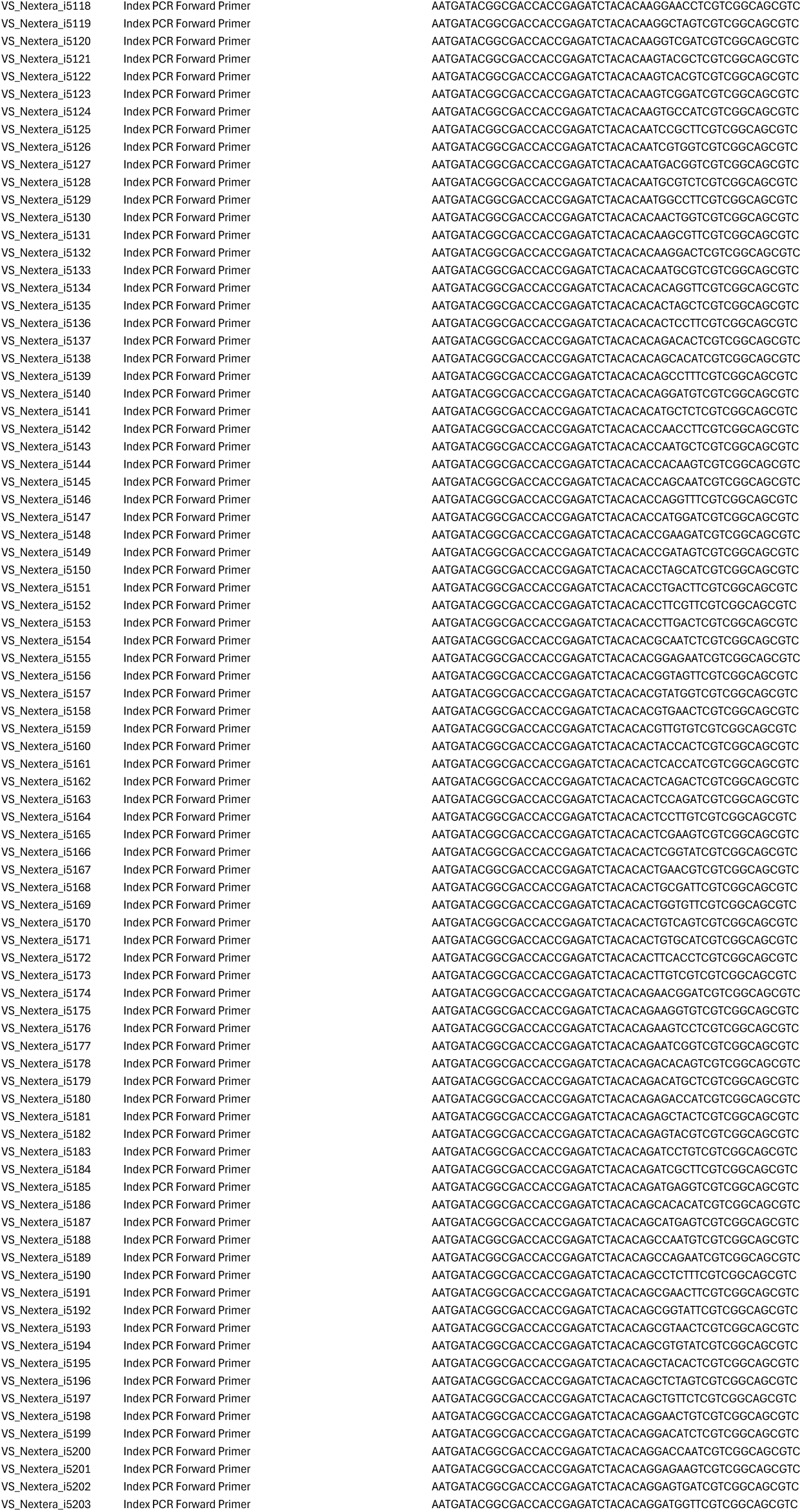
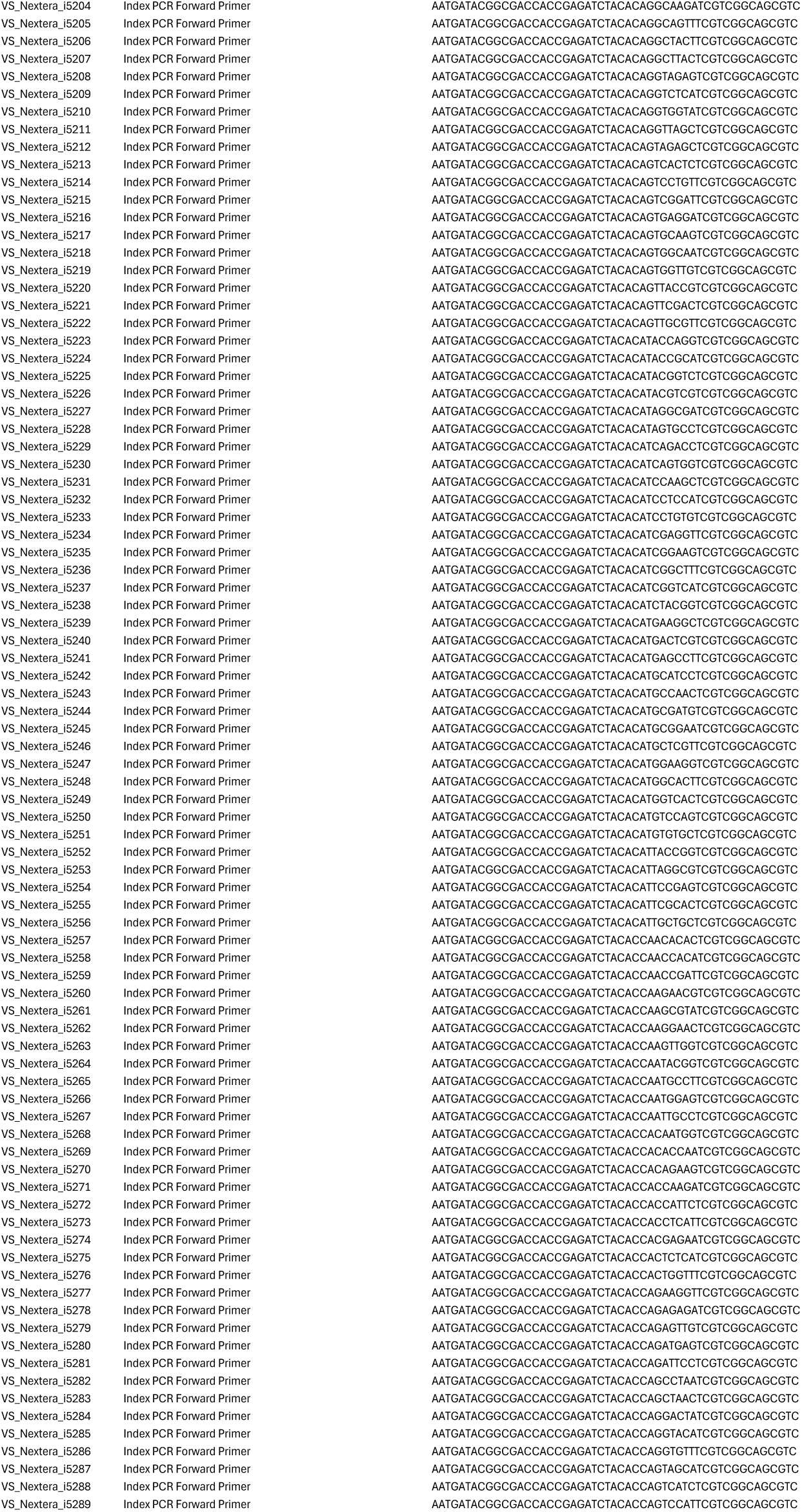
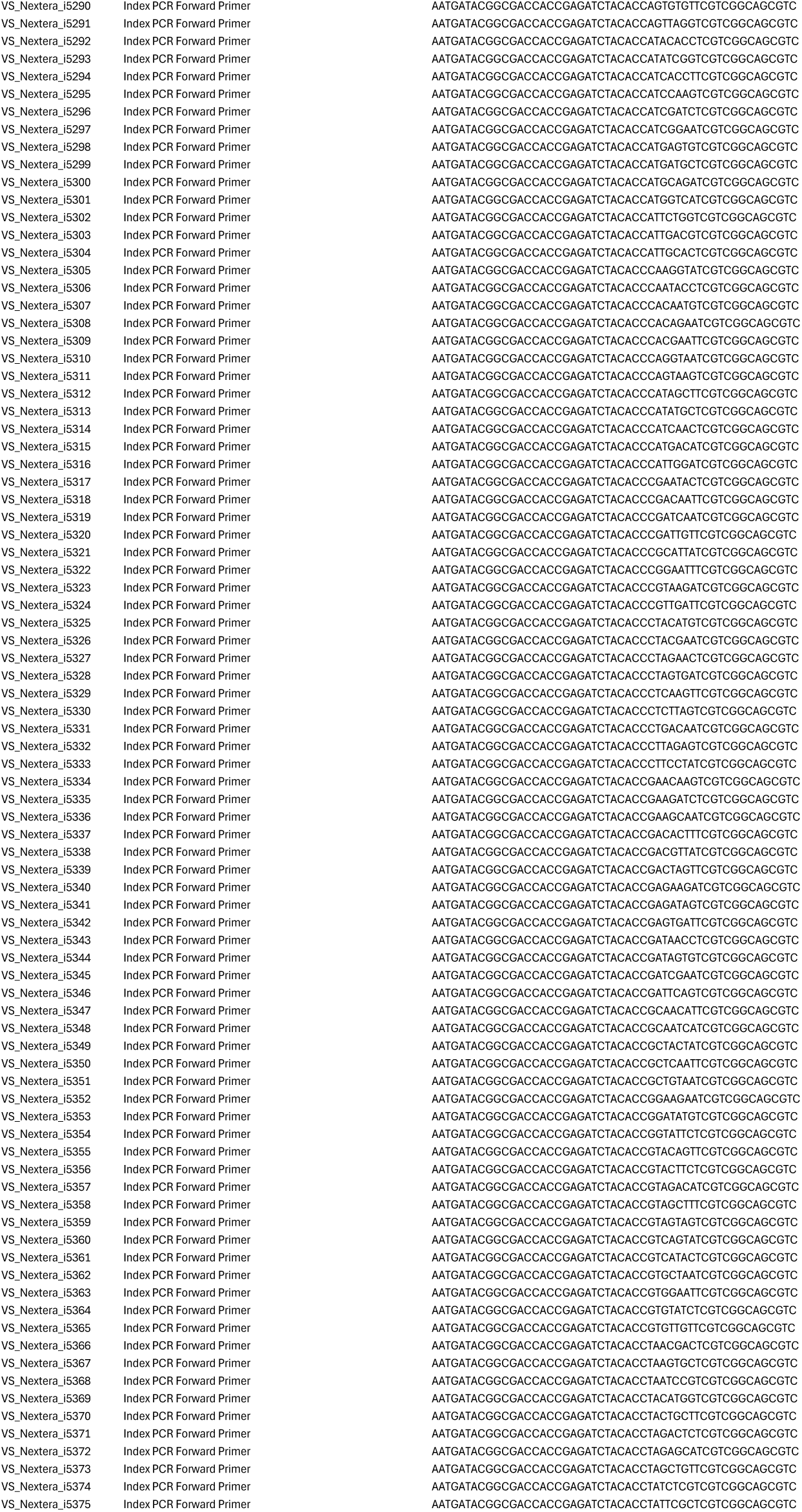
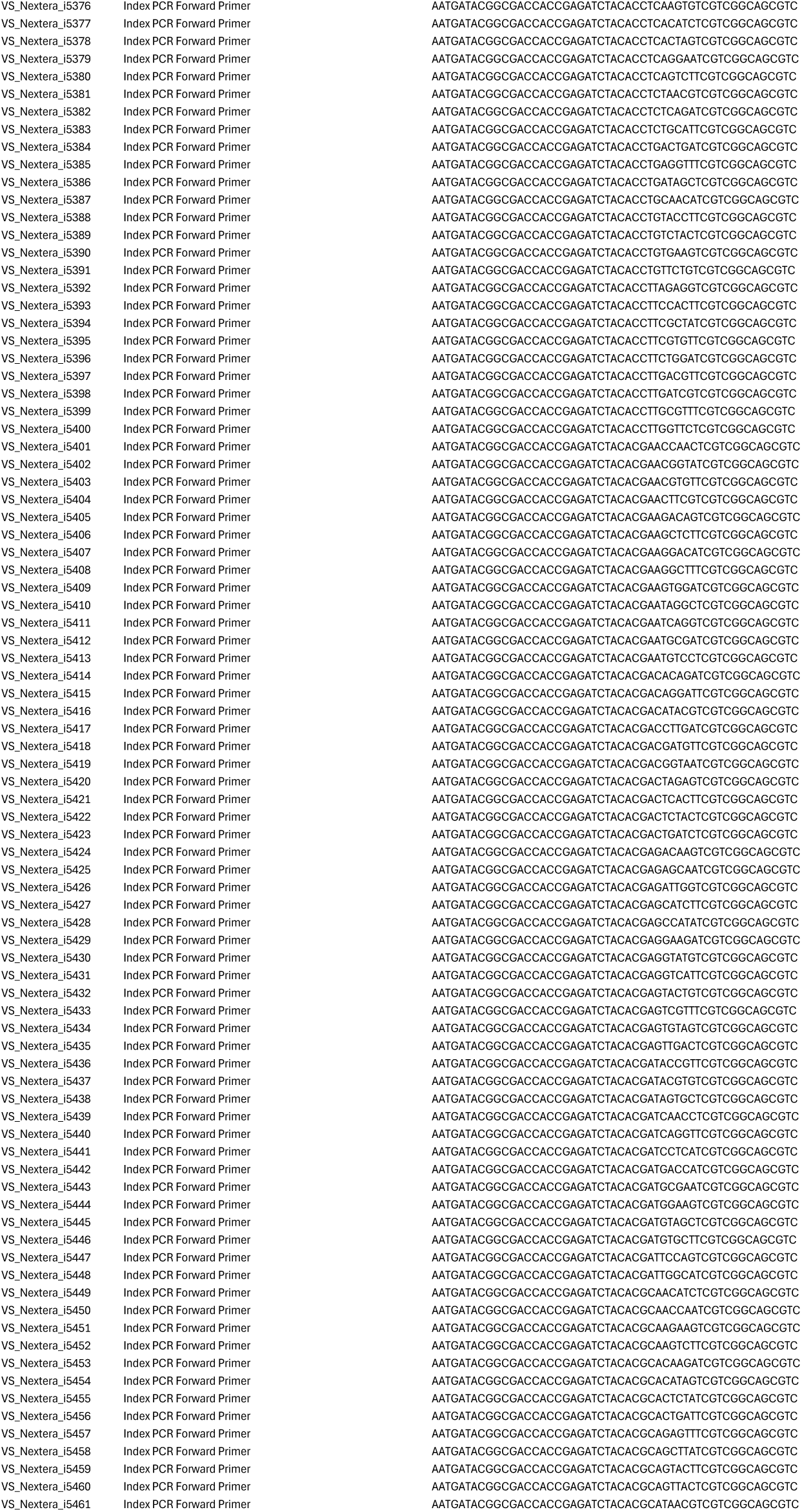
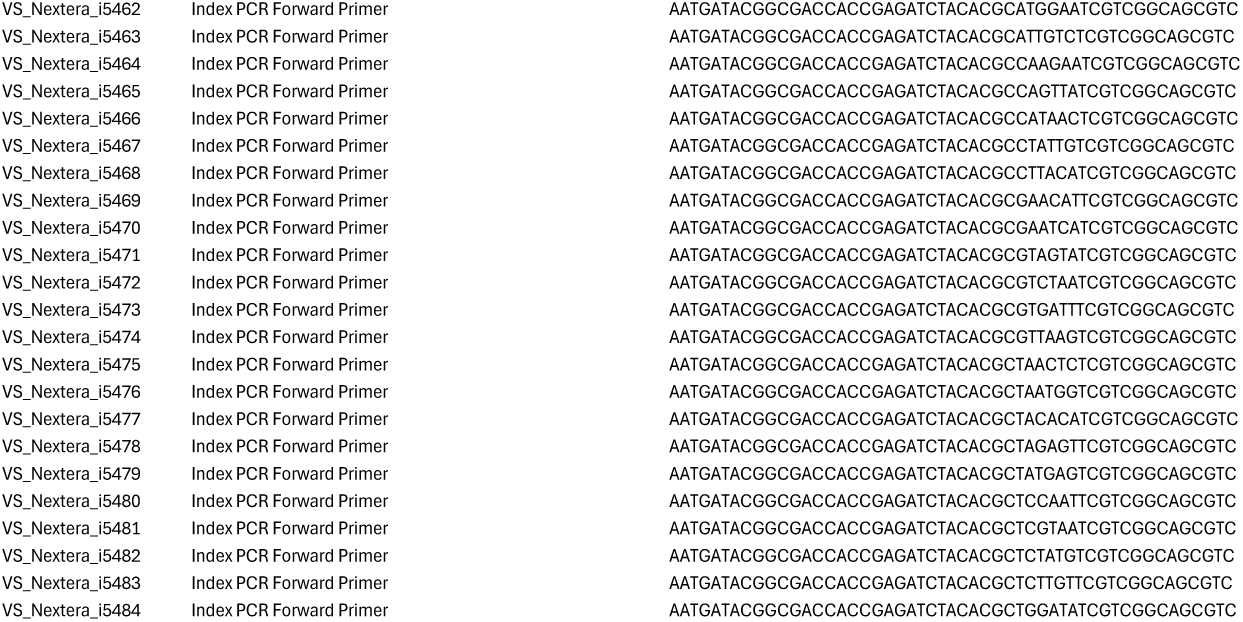
List of all DNA oligos and PCR primers used in the present study.

